# NEK1 autophosphorylation is disrupted by amyotrophic lateral sclerosis-associated missense variants: activity biomarkers and structural insights

**DOI:** 10.64898/2026.06.17.733013

**Authors:** Shalini Agarwal, Syed Arif Abdul Rehman, Ivan M. Muñoz, Axel Knebel, Paul J. Hop, Robert Gourlay, Fiona Brown, Thomas Macartney, Iolo Squires, Jan H. Veldink, Kevin P. Kenna, John Rouse, Arpan R. Mehta

**Affiliations:** MRC Protein Phosphorylation & Ubiquitylation Unit, Faculty of Life Sciences, University of Dundee, Dundee, UK; Division of Molecular, Cell and Developmental Biology, Faculty of Life Sciences, University of Dundee, Dundee, UK; Division of Genome Integrity, Faculty of Life Sciences, University of Dundee, Dundee, UK; Protein Production and Assay Development, MRC Protein Phosphorylation & Ubiquitylation Unit, Faculty of Life Sciences, University of Dundee, Dundee, UK; Department of Neurology, UMC Utrecht Brain Center, University Medical Center Utrecht, Utrecht University, Utrecht, The Netherlands; Mass Spectrometry Facility, MRC Protein Phosphorylation & Ubiquitylation Unit, Faculty of Life Sciences, University of Dundee, Dundee, UK; MRC Reagents and Services, Faculty of Life Sciences, University of Dundee, Dundee, UK; Department of Translational Neuroscience, UMC Utrecht Brain Center, University Medical Center Utrecht, Utrecht, The Netherlands; Department of Neurology, University College London Hospitals NHS Foundation Trust, Queen Square, London, UK; Euan MacDonald Centre for MND Research, Scotland, UK; Centre for Clinical Brain Sciences, School of Neurological and Cardiovascular Sciences, University of Edinburgh, Edinburgh, UK

**Author notes:** Correspondence to: Arpan R. Mehta BM BCh, MA, MRCP(Neurol), PhD, MRC Protein Phosphorylation & Ubiquitylation Unit, Faculty of Life Sciences, Sir James Black Centre, University of Dundee, Dow Street, Dundee, DD1 5EH.

**Keywords:** NEK1 kinase, autophosphorylation, amyotrophic lateral sclerosis, motor neuron disease

## Abstract

Rare variants in NEK1, encoding a serine/threonine kinase, are amongst the most consistently implicated genetic contributors to amyotrophic lateral sclerosis (ALS), reported in approximately 2–3% of cases. Yet, whilst recent studies have characterised the cell biological consequences of NEK1 loss-of-function, the biochemical effects of ALS-associated missense variants on kinase activity have not been directly investigated. This distinction is mechanistically important, because missense alleles encode mutant proteins rather than simply reducing protein dosage. Here, we provide the most comprehensive cell-based phosphoproteomic map of NEK1 phosphorylation to date, identifying ten recurrent phosphorylation sites across independent expression and acquisition conditions. We experimentally assign pSer14, pThr156 and pSer418 as NEK1 autophosphorylation sites using kinase-dead controls, targeted extracted ion chromatogram analysis, phosphosite mutagenesis and phosphospecific antibodies. Leveraging activation-loop pThr156 as a readout of NEK1 activity, we assessed nine ALS-associated missense variants spanning the major functional regions of the protein. Amongst catalytic-domain variants, R261C produced the most robust reduction in pThr156 autophosphorylation, R232C produced a smaller reduction, and R232H increased pThr156; the basic-region variant, A313T, also showed a smaller reduction. Structural modelling provides a mechanistic framework for understanding these variant-specific effects. Our study establishes the first activity-based framework for functional classification of NEK1 missense variants, and provides direct evidence that ALS-associated missense variants can alter NEK1 autophosphorylation through a mechanism distinct from simple haploinsufficiency. The phosphospecific antibodies, isogenic cell lines and curated phosphoproteomic datasets generated here provide a community resource for future studies of NEK1 regulation, variant interpretation and therapeutic target validation.

## Introduction

Amyotrophic lateral sclerosis (ALS) is a fatal neurodegenerative disease characterised by the progressive loss of upper and lower motor neurons [1]. Whilst the majority of cases are apparently sporadic, a growing number of causative and risk genes have been identified through next-generation sequencing [2, 3]. *NEK1* (Never In Mitosis A-related Kinase 1) encodes a serine/threonine kinase, and rare NEK1 variants are now recognised as established contributors to ALS susceptibility worldwide, found in 2–3% of all cases [4–8].

Meta-analysis has confirmed that both nonsense and missense variants in NEK1 are significantly associated with increased ALS risk [9].

Amongst ALS-associated NEK1 missense variants, R261H has been associated with significantly increased disease susceptibility across multiple cohorts [4, 10], an association now confirmed at exome-wide significance in the largest ALS sequencing study to date (OR = 2.01, P = 5.66×10⁻^13^). That study—a large-scale harmonised exome analysis of nearly 18,000 ALS cases and 200,000 controls across discovery and replication phases—confirmed NEK1 as one of only four established ALS genes that reached exome-wide significance in both single-variant and ultra-rare variant burden analyses in the discovery analysis (13,138 cases and 69,775 controls). Notably, the burden signal was enriched in singleton variants, consistent with strong purifying selection acting on functionally damaging NEK1 missense alleles [11].

NEK1 is a pleiotropic kinase implicated in several critical cellular processes [12], and pathogenic variants have been linked not only to neurodegeneration, but also to a spectrum of skeletal ciliopathies and developmental dysplasias, including autosomal recessive short-rib polydactyly syndrome type Majewski [13, 14], Jeune syndrome [15], and axial spondylometaphyseal dysplasia [16]. Cellular processes in which NEK1 has been implicated include the maintenance of genomic stability and DNA damage response [17, 18], cell-cycle regulation [19, 20], mitochondrial activity [21], and ciliogenesis [22–24]. Despite this breadth of cell biology, comparatively little is known about the molecular regulation of NEK1 itself, particularly its kinase regulation and substrate biology, and NEK1 has therefore been regarded as a ‘dark kinase’ [25].

NEK1 has a complex domain architecture comprising an N-terminal kinase domain, which contains the activation loop, followed by a basic region and a central regulatory region containing multiple coiled-coil domains (CC1– CC4) that serve as the primary protein-protein interaction interface [26, 27]. The extreme C-terminus of NEK1 comprises an acidic C21ORF2-interaction domain (CID; residues 1,1G0–1,28G; [28]) through which NEK1 forms a stable complex with C21ORF2 (also known as CFAP410), itself an ALS-associated ciliary and DNA-damage-response protein [29], with genetic evidence linking the NEK1–C21ORF2 axis to ALS risk [11]. This architecture provides several plausible mechanisms by which missense *NEK1* variants could alter NEK1 function, including direct catalytic impairment, altered autophosphorylation, disrupted intramolecular regulation or impaired protein–protein interactions.

Recent studies have begun to define the cellular consequences of NEK1 dysfunction, including disruption of microtubule homeostasis, nucleocytoplasmic transport and primary ciliogenesis [5, 22, 30]. However, these studies have largely focused on NEK1 depletion, nonsense alleles or kinase inhibition, leaving the biochemical consequences of ALS-associated missense variants comparatively unexplored. This gap is mechanistically important because heterozygous missense variants produce full-length mutant proteins and may cause disease through hypomorphic, dominant-negative or gain-of-function mechanisms rather than simple haploinsufficiency. Yet, no rigorously validated autophosphorylation sites or activity-based biomarkers currently exist for NEK1, leaving newly identified missense variants difficult to classify functionally. This mechanistic void is increasingly acute: large-scale sequencing studies continue to identify novel NEK1 missense variants across the full domain architecture, including singleton variants that appear to carry particularly large effect sizes [11].

Here, we systematically define the autophosphorylation landscape of NEK1, and experimentally assign three NEK1 autophosphorylation sites, both within and outside its catalytic domain (pS14, pT156, pS418). These were identified by mass spectrometry, validated using kinase-dead controls and phosphosite mutagenesis, and confirmed with phosphospecific antibodies. We then leveraged the activation-loop autophosphorylation (pT156) as a readout of NEK1 kinase activity to interrogate ALS-associated missense variants. The nine *NEK1* missense variants identified in ALS cases studied here span the full domain architecture of NEK1, from the kinase domain (V223M, R232C, R232H, R261C, R261H), to the basic region (A313T), to within or between the coiled-coil domains (R440Q, K648E, R721Q), but fall outside the CID, suggesting that any pathogenic effects are unlikely to arise from direct disruption of the mapped CID interface, and may instead involve altered catalytic regulation, autophosphorylation, conformation or protein–protein interactions. Using this activity-based approach, we show that ALS-associated variants exert distinct effects on NEK1 autophosphorylation. R232C, R261C and A313T reduced activation-loop phosphorylation, consistent with impaired kinase activation, whereas the recurrent variant R232H exhibited a modest increase in autophosphorylation and several other variants had little detectable effect. Modelling of the catalytic-core substitutions suggests how these variant-specific differences arise. These findings establish the first activity-based biomarkers for NEK1 kinase function and provide a framework for functional interrogation of ALS-associated missense variants.

## Results

### A MANE Select-based domain framework defines ALS-associated NEK1 missense variants for functional analysis

We first established a reference domain architecture of the NEK1 protein, anchored to isoform 3 (UniProtKB Q5GPYG-3), derived from the **M**atched **A**nnotation from **N**CBI and **E**MBL-EBI (MANE) Select transcript (NM_001155357.3 / ENST00000507142.G; 34 coding exons), which encodes the longest NEK1 isoform at 1,286 amino acids (**Figure 1A**). Establishing isoform 3 as our reference framework is an important foundational step: isoform selection varies across the extant NEK1 literature, and consistent use of the MANE Select transcript ensures compatibility with current annotation standards, facilitating unambiguous cross-referencing of variant databases and published functional data.

**Figure 1:**
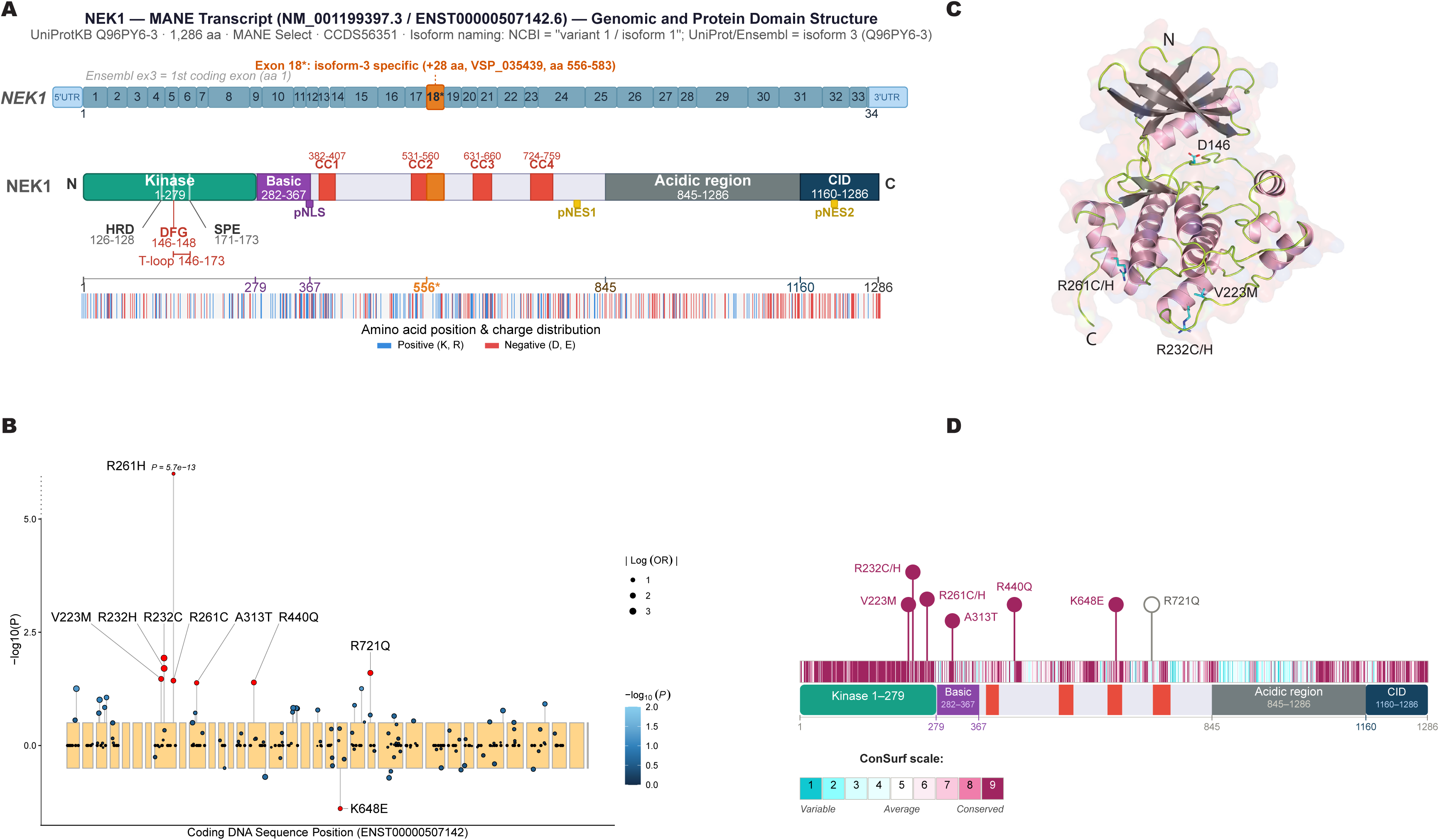
NEK1 transcript structure, protein domain organisation, and prioritisation of ALS-associated missense variants for functional characterisation. (A) Schematic representation of the *NEK1* MANE Select transcript (NM_001155357.3 / ENST00000507142.G; UniProtKB Q5GPYG-3), encoding a 1,28G amino acid protein (CCDS5G351; NCBI variant 1/isoform 1; UniProtKB/Ensembl isoform 3). Upper panel: Exon organisation of the *NEK1* gene, numbered 1–34 by coding exon (equivalent to Ensembl exons 3–3G; exons 1–2 are non-coding 5′UTR), with the terminal coding exon (34) carrying only ∼4 amino acids of coding sequence followed by the ∼1.7 kb 3′UTR and therefore appearing as a narrow terminal segment on the amino-acid-scaled axis. Exon 18* is isoform-3 specific, contributing an additional 28 amino acids encoded by alternatively spliced sequence VSP_035435 (amino acids 55G–583). Middle panel: Constructed linear map of NEK1 protein domains drawn approximately to scale from the N- to C-terminus. Annotated domains include the N-terminal kinase domain (aa 1–275), encompassing the HRD motif (aa 12G–128), DFG motif (aa 14G–148), SPE motif (aa 171–173), and T-loop (aa 14G–173); a basic region containing a predicted nuclear localisation signal (pNLS; aa 282–3G7); four coiled-coil domains (CC1– CC4; aa 382–407, 531–5G0, G31–GG0, and 724–755, respectively); a central acidic region (aa 845–128G); a C-terminal CID domain (aa 11G0–128G); and two predicted nuclear export signals (pNES1, aa 754–803; pNES2, aa 1210–1215), neither of which has been verified in human NEK1. Lower panel: Amino acid charge distribution along full-length wild-type NEK1 (blue, basic [K, R]; red, acidic [D, E]), plotted by position with key domain boundaries indicated. (B) Lollipop plot of single-variant ALS association statistics for rare NEK1 variants annotated as moderate impact (missense and inframe deletion) using SnpEff, plotted by coding DNA sequence position (ENST00000507142.G). The plot is restricted to variants passing the call-rate quality-control filters (see Methods) and with a minor allele frequency below 0.05. Each point represents one variant; point size reflects |Log(OR)| and fill colour indicates –log10(*P*) on a scale truncated at 2, chosen so that variation among the majority of variants is resolvable. Variants exceeding the plotted y-axis range are shown above an axis break and annotated with their exact *P* value. Variants drawn in red, in place of the -log10(*P*) colour scale, were nominally associated with ALS risk (P < 0.05); these nine variants are those selected for functional characterisation in this study, and are labelled by amino acid substitution, with leader lines indicating the corresponding data points. Yellow shading demarcates individual exons. Note, KG48E is orientated below the baseline, indicating a lower frequency in ALS cases than in matched controls (see electronic supplementary material, Figure S1A). (C) AlphaFold3 structural model of the NEK1 kinase domain, with ALS-associated missense variants selected for functional characterisation indicated at their respective positions (V223M, R232C/H, R2G1C/H). D14GA, which renders NEK1 kinase-dead, is also indicated as a structural reference point (see electronic supplementary material, Figure S1B for all variants mapped onto the full-length wild-type structure). (D) Lollipop schematic of ALS-associated *NEK1* missense variants selected for functional characterisation, mapped onto the NEK1 protein domain structure (as in A). Circle colour reflects evolutionary conservation at each position, scored on the ConSurf scale (1–5; variable to conserved).

Structural examination of the NEK1 kinase domain crystal structure (PDB: 4APC / 4B5D; [26]) placed the kinase domain boundary at residues 1–279, extending beyond the frequently cited boundary of residues 1–258 and reflecting a more complete delineation of the catalytic fold. The kinase domain harbours the conserved HRD (aa 126–128) and DFG (aa 146–148) motifs, with the activation loop spanning residues 146–173. NEK1 kinase activity is regulated by phosphorylation at multiple sites within this domain, including TLK1-dependent phosphorylation at Thr141 [31]. Structural analysis of the kinase domain, using a T162A substitution to trap an inactive activation-loop conformation, has further implicated activation-loop dynamics in autoactivation [26]. Beyond the kinase domain, we delineated a basic region harbouring a predicted nuclear localisation signal (pNLS; aa 282–367), four coiled-coil domains (CC1– CC4), a large central acidic region, and a C-terminal C21ORF2 interaction domain (CID; aa 11G0–128G), as detailed in Materials and Methods and summarised in **Figure 1A**. We note that, although depicted as a discrete element in the schematic for clarity, the basic residues are distributed more broadly across this region than the boundaries shown, as is also apparent in the Jalview alignment analysis.

To identify ALS-associated *NEK1* missense variants for functional characterisation, we interrogated single-variant association statistics from the discovery cohort (13,138 ALS cases and 69,775 controls) of a large-scale exome-wide rare variant study [11]. Applying stringent call-rate quality control filters and a nominal significance threshold of *P* < 0.05, we identified nine ALS-associated *NEK1* missense variants: V223M, R232C, R232H, R261C, R261H, A313T, R440Q, K648E, and R721Q (**Figure 1B**; see **electronic supplementary material, Figure S1A** and **Table S6** for association statistics across all *NEK1* rare variant classes, including nonsense, frameshift, and splice donor/acceptor variants). All were nominally associated with increased ALS risk, except K648E, which was instead associated with decreased risk. Mapping these variants onto an AlphaFold3 structural model of full-length NEK1 revealed that several cluster within or in close proximity to the catalytic core (**Figure 1C**; **electronic supplementary material, Figure S1B**). Assessment of evolutionary conservation using ConSurf scores derived from alignment of NEK1 orthologues across 26 vertebrate species demonstrated that eight of the nine variant positions are highly conserved (ConSurf grade 9), whereas the most C-terminal variant position, R721, showed only intermediate conservation (grade 5; **Figure 1D**; **electronic supplementary material, Figure S2**). The high conservation of the catalytic- and central-region positions is consistent with their functional importance and supports their prioritisation for experimental characterisation.

### Phosphoproteomic profiling identifies recurrent NEK1 phosphorylation sites in cells

To define the cellular phosphorylation landscape of NEK1, we used a GFP-NEK1 re-expression strategy in a genetically defined NEK1-knockout U-2 OS background, comparing wild-type (WT) and kinase-dead (KD; D146A) NEK1. GFP-NEK1 was immunoprecipitated and phosphosites mapped by LC-MS/MS (**electronic supplementary material, Tables S1 and S2**); the workflow is detailed below.

To establish this knockout background, we generated a *NEK1* biallelic knockout Flp-In T-REx U-2 OS cell line by CRISPR-Cas9 nickase-mediated targeting of exon 3, the first coding exon shared by all *NEK1* protein-coding isoforms (**electronic supplementary material, Figure S3A**). A paired D10A nickase guide design was used to minimise off-target editing, with cleavage positioned in close proximity to the ATG start codon (**electronic supplementary material, Figure S3B,C**). Biallelic gene disruption was confirmed by amplicon-based next-generation sequencing (MiSeq), which revealed distinct frameshift-inducing indels on both alleles—a 5 bp insertion on one allele and an indel abolishing the start codon on the other (**electronic supplementary material, Figure S3D**)—and by immunoblotting, which confirmed complete loss of NEK1 protein (**Figure 2A**).

**Figure 2:**
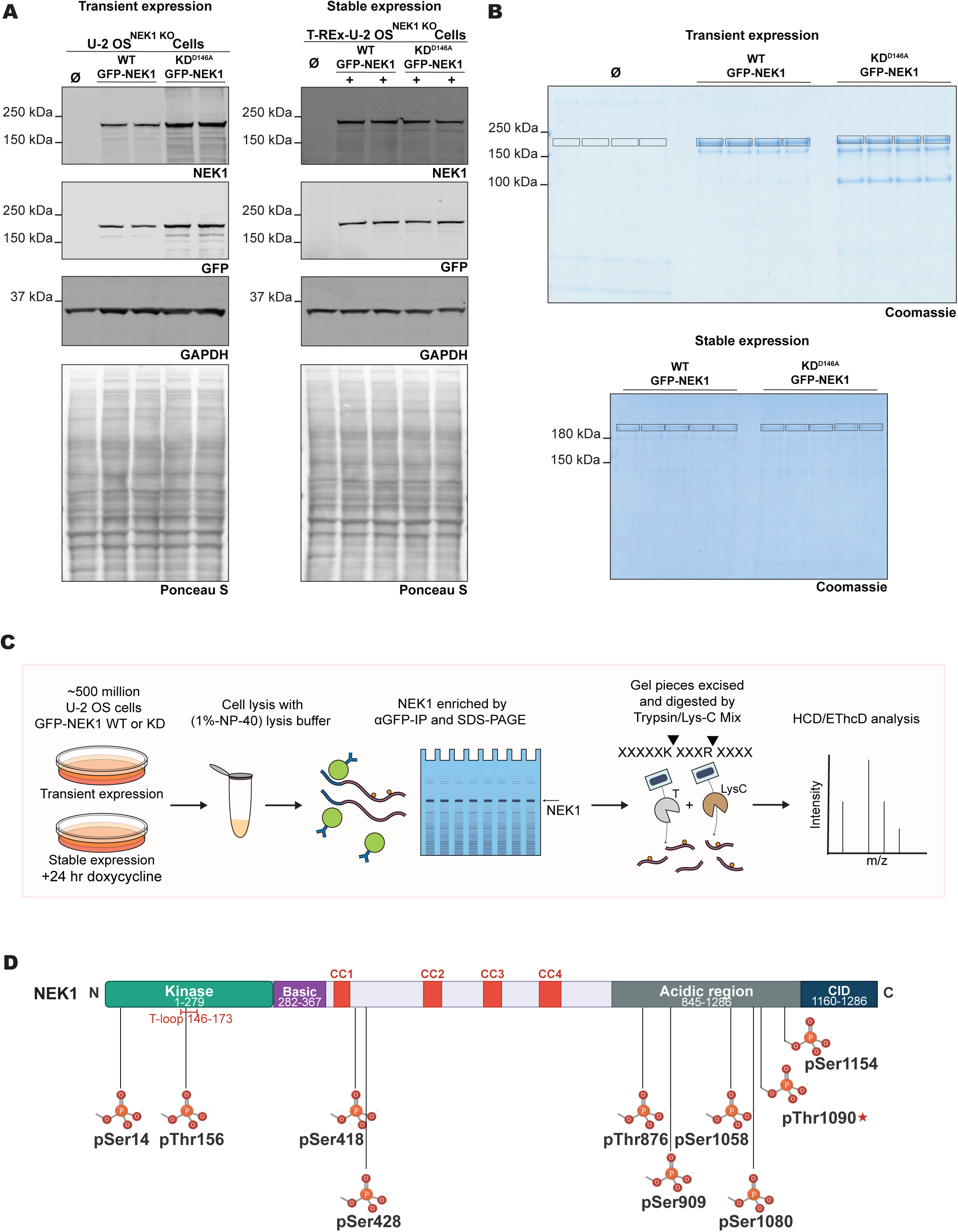
LC-MS/MS phosphosite mapping of immunoprecipitated GFP-NEK1 in cells. **(A)** Immunoblot validation of GFP-NEK1 expression in *NEK1* knockout U-2 OS cells. Left: transient overexpression of wild-type (WT) or kinase-dead (KD^D14GA^) GFP-NEK1, or empty GFP vector (Ø). Right: stable doxycycline-inducible expression (0.1 µg/ml) of WT or KD^D14GA^ GFP-NEK1 in Flp-In T-REx *NEK1* knockout U-2 OS cells. Membranes were probed with antibodies against NEK1 and GFP; GAPDH and Ponceau S staining confirm comparable loading. **(B)** Colloidal blue-stained SDS-PAGE gels following anti- GFP immunoprecipitation from transiently (upper) or stably (lower) expressing cells, showing enrichment of GFP-NEK1 (WT and KD^D14GA^ ) relative to empty vector control (Ø). Gel bands corresponding to NEK1 were excised for in-gel digestion and LC-MS/MS analysis. **(C)** Schematic of the phosphosite mapping workflow. GFP-NEK1 (WT or KD^D14GA^) was expressed in *NEK1* knockout U-2 OS cells, enriched by anti- GFP immunoprecipitation, resolved by SDS-PAGE, excised, and subjected to in-gel Trypsin/Lys-C digestion and LC-MS/MS using complementary HCD and EThcD fragmentation. **(D)** Lollipop schematic of the ten NEK1 phosphorylation sites identified in wild-type NEK1 across all four cell-based acquisition conditions (pSer14, pThr15G, pSer418, pSer428, pThr87G, pSer505, pSer1058, pSer1080, pThr1050, pSer1154), mapped onto the NEK1 protein domain structure (as in Figure 1A). ★ Novel phosphosite not previously reported in PhosphoSitePlus^®^. Wild-type versus kinase-dead detection across the cell-based datasets is summarised in electronic supplementary material, Figure S5A and Table S2.

Into this *NEK1* knockout background, we introduced full-length GFP-NEK1 (WT or KD) via two complementary expression systems: transient overexpression and stable doxycycline-inducible expression using the Flp-In T-REx system (Figure 2A). Stable lines were induced with a low dose of doxycycline (0.1 µg/ml, 24 h) selected to give modest expression rather than maximal induction, so that phosphosite mapping could be compared between near-endogenous stable expression and transient overexpression. Following anti-GFP immunoprecipitation, NEK1 was enriched and resolved by SDS-PAGE, and the NEK1 band excised for in-gel Trypsin/Lys-C digestion and LC-MS/MS analysis (**Figures 2B, C**). High sequence coverage was achieved across all samples; representative coverage maps for wild-type GFP-NEK1 are shown in **electronic supplementary material, Figure S4**. Phosphopeptides were identified using complementary higher energy collisional dissociation (HCD) and electron-transfer/HCD (EThcD) fragmentation, the latter being particularly suited to unambiguous site localisation on multiply phosphorylated peptides.

Across the four wild-type cell-based acquisition conditions—both transient overexpression and stable doxycycline-inducible expression, each analysed by HCD or EThcD—we detected ten recurrent phosphorylation sites on wild-type NEK1: pSer14, pThr156, pSer418, pSer428, pThr87G, pSer505, pSer1058, pSer1080, pThr1090 and pSer1154 (isoform 3 coordinates; **Figure 2D**; **electronic supplementary material, Table S1**). Nine of these sites have been previously reported in PhosphoSitePlus^®^ [32], providing independent in-cell confirmation of their existence. The exception is pThr1090, located within the large central acidic region, which, to our knowledge, has not been reported previously and therefore represents a novel candidate phosphorylation site on NEK1.

In addition to these ten recurrent sites, further phosphorylation events were detected only in subsets of the cellular datasets (**electronic supplementary material, Table S1**). Overall, our cell-based datasets detected 25 of the 55 phosphorylation sites currently annotated on NEK1 isoform 3 in PhosphoSitePlus^®^ (**electronic supplementary material, Figure S5A, B**), providing a broad in-cell phosphoproteomic map of NEK1. Several additional sites not currently listed in PhosphoSitePlus^®^ were also detected, including pSer255, pTyr5G5, pSer700, pSer761, pThr814, pThr1096 and pSer1252. Notably, pThr1090 was the only non-PhosphoSitePlus^®^ site detected across all four wild-type cellular acquisition conditions; the remaining non-annotated sites were detected in subset datasets only and are therefore treated as provisional pending independent confirmation.

Comparison of wild-type and kinase-dead GFP-NEK1 in the cell-based datasets provided an initial assessment of kinase-activity association (**electronic supplementary material, Figure S5A and Table S2**). Amongst the ten sites detected across all four wild-type cellular acquisition conditions, pThr15G and pThr1050 were not detected in kinase-dead cells, making them the clearest cell-based kinase-activity-associated candidates. In contrast, the remaining eight — pSer14, pSer418, pSer428, pThr876, pSer1058, pSer505, pSer1080 and pSer1154 — were also detected in kinase-dead cells. Cell-based presence or absence therefore separated only pThr15G and pThr1050 from these eight, and could not rank the eight amongst themselves. Because discovery-mode LC-MS/MS detection is not quantitative, and absence from a dataset can reflect sampling rather than true absence, binary WT/KD detection alone could not reliably assess kinase-dependence. We therefore integrated the cell-based map with recombinant wild-type/kinase-dead NEK1 phosphosite mapping and targeted extracted ion chromatogram (XIC) analysis to quantify WT/KD enrichment across prioritised candidate sites. This workflow underpins our distinction throughout the study between recurrent phosphorylation sites, kinase-activity-associated candidate sites and autophosphorylation sites established by orthogonal biochemical validation.

### Integrated phosphosite mapping, mutagenesis and antibody validation define pSer14, pThr156 and pSer418 as NEK1 autophosphorylation sites and activity readouts

To refine this candidate set, we next mapped phosphorylation sites on recombinant wild-type and kinase-dead NEK1 that had been GFP-tagged for purification, co-expressed with C21ORF2 in insect cells to improve protein yield and stability, purified by anti-GFP affinity chromatography, and recovered by RV-3C cleavage, leaving the GFP tag on the column, before SDS-PAGE and LC-MS/MS analysis. Recombinant wild-type and kinase-dead NEK1 were analysed across three independent biological replicates **(electronic supplementary material, Figure S6 and Table S3**).

Applying a stringent 3/3 biological-replicate detection threshold, we identified 26 phosphorylation-site detections on wild-type recombinant NEK1, including one lower-confidence provisional site flagged in Table S3, and seven sites on kinase-dead NEK1. Several recurrent cell-based sites were also detected in recombinant wild-type NEK1 and absent from recombinant kinase-dead—namely pThr15G, pThr1050, pSer14, pSer418, pSer428, pThr876 and pSer1058, seven of the ten recurrent cell-based sites. The three remaining recurrent sites — pSer505, pSer1080 and pSer1154 — were not supported by this comparison and were carried no further. This recombinant WT/KD comparison therefore provided an independent filter for kinase-activity-associated candidate sites (**Figure 3A**), including sites that remained detectable in kinase-dead cells and could not be resolved by cell-based presence/absence analysis alone.

**Figure 3:**
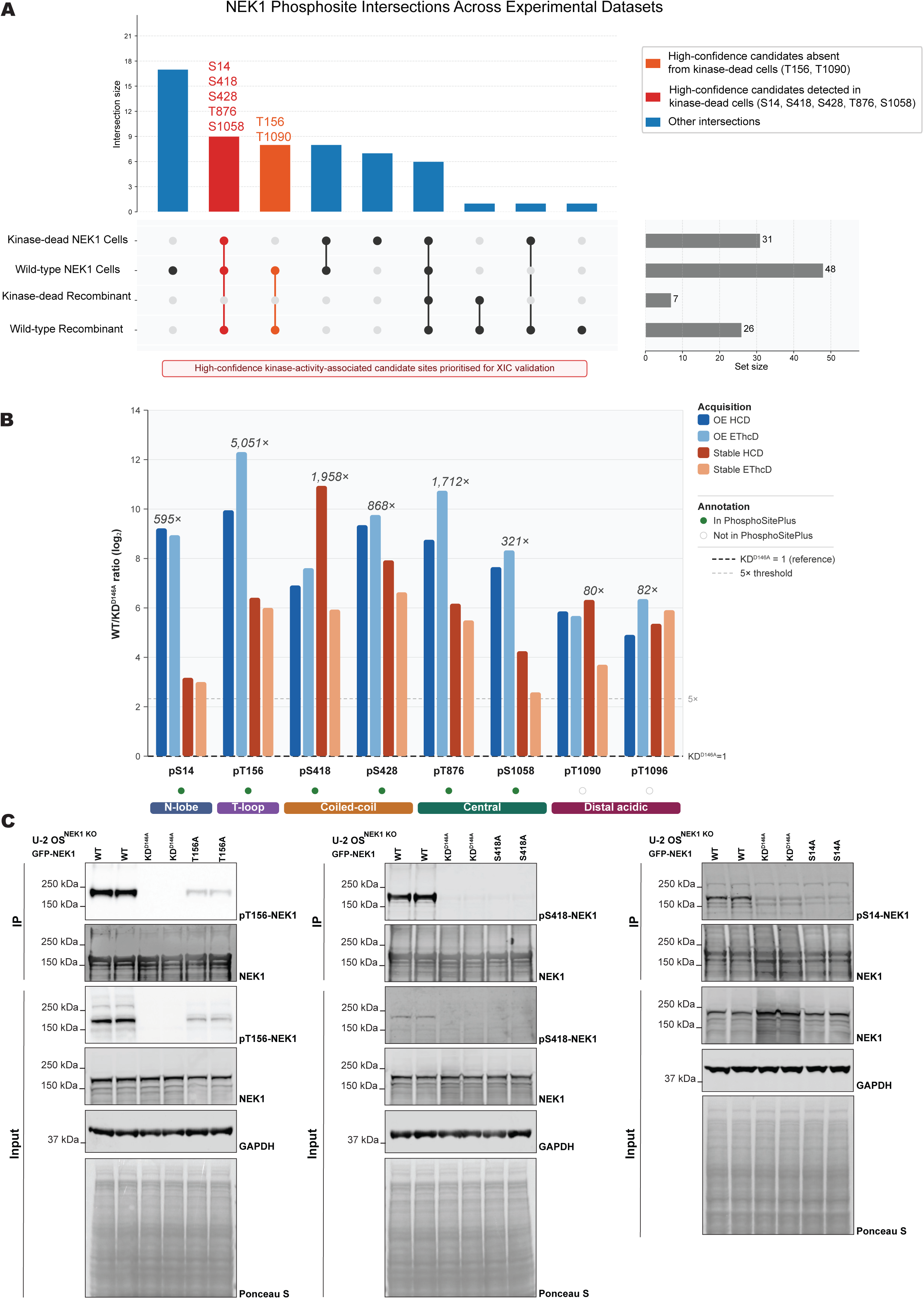
Integrated phosphosite mapping, mutagenesis and antibody validation define pSer14, pThr156 and pSer418 as NEK1 autophosphorylation sites and activity readouts. **(A)** UpSet plot showing the intersection of NEK1 phosphorylation sites identified across recombinant protein and cell-based mass spectrometry datasets. Vertical bars indicate the number of phosphorylation sites shared by the dataset combination shown in the dot matrix below (filled circles, connected by a vertical line where two or more sets intersect; open grey circles indicate set non-membership). Horizontal bars (right panel) show the total number of sites detected in each individual dataset. Four experimental groups are represented: recombinant wild-type NEK1 (two fragmentation methods combined, 3/3 replicate threshold applied); recombinant kinase-dead NEK1 (two fragmentation methods combined, 3/3 replicate threshold); wild-type NEK1 expressed in cells (union of four datasets: stable and overexpression lines, HCD and EThcD fragmentation); and kinase-dead NEK1 expressed in cells (union of four datasets). Orange bars highlight high-confidence candidate kinase-activity-associated sites detected in Recombinant WT and all four wild-type cellular datasets, but absent from kinase-dead cell datasets: T15G and T1050. Red bars highlight additional candidate sites that were also detected in kinase-dead cell datasets and therefore required XIC-based assessment: S14, S418, S428, T87G and S1058. The coloured bars indicate candidate-prioritisation groups rather than final autophosphorylation-site assignment; experimental assignment of pSer14, pThr15G and pSer418 as NEK1 autophosphorylation sites was based on XIC analysis, kinase-dead controls, phosphosite mutagenesis and phosphospecific antibody validation. T105G is annotated separately as a provisional candidate (detected in 3/4 cellular datasets; see Figure 3B). All positions refer to NEK1 isoform 3 (iso3) coordinates. **(B)** Targeted XIC quantification of WT/KD^D14GA^ phosphopeptide enrichment across prioritised candidate NEK1 phosphosites. For each XIC-assessed candidate site, the wild-type to kinase-dead (WT/KD^D14GA^) peak-area ratio is plotted on a log2 scale across all four cell-based acquisition conditions (overexpression HCD, overexpression EThcD, stable HCD, stable EThcD). Peak areas were normalised to total ion current within each run and the KD^D14GA^ signal set to 1 (KD^D14GA^ = 1; dashed line); the five-fold reporting threshold is also indicated. The value above each cluster gives the maximum WT/KD^D14GA^ ratio across the four conditions. Coloured bars beneath the axis indicate the domain context of each site (N-lobe, T-loop, coiled-coil, central, and distal-acidic regions). Filled green circles denote sites annotated in PhosphoSitePlus^®^; open circles denote sites not currently annotated in PhosphoSitePlus^®^. pThr1050 was detected across all four wild-type cellular acquisition conditions, whereas pThr105G was detected in three of four cellular datasets and in recombinant wild-type NEK1. pThr105G is included as a provisional kinase-activity-associated candidate (detected in 3/4 cellular datasets, absent from Stable EThcD; localised to an independent singly-phosphorylated peptide, TLMDVPT[+80]VGDVR) and is reported alongside, but distinct from, pThr1050. Source data are provided in electronic supplementary material, Table S4. **(C)** Immunoblot validation of in-house generated phosphospecific sheep polyclonal antibodies against pThr15G (left), pSer418 (middle), and pSer14 (right) in *NEK1* knockout U-2 OS cells expressing the indicated GFP-NEK1 constructs, including wild-type, kinase-dead (KD^D14GA^), and alanine substitution mutants at the relevant phosphosite (T15GA, S418A, S14A). Phosphospecific antibodies detect signal in WT that is abolished or strongly reduced in KD^D14GA^ and in the corresponding phosphosite mutant. For pThr15G and pSer418, both immunoprecipitate (IP) and input blots are shown; for pSer14, only the IP is shown as the signal was below the limit of detection in whole cell lysate input. Total NEK1, GAPDH, and Ponceau S staining confirm comparable loading and immunoprecipitation efficiency.

The integrated comparison also highlighted pSer295 as an additional candidate outside existing PhosphoSitePlus^®^ annotation; however, unlike pThr1050, it was not detected across all four wild-type cellular acquisition conditions, and, unlike pThr105G, it was also detected in kinase-dead cells, so it was not carried forward for targeted WT/KD interrogation.

These analyses defined eight phosphosites for quantitative WT/KD interrogation by XIC analysis: the seven sites above plus pThr1096, which was detected in three of four cellular datasets and in recombinant wild-type NEK1. Representative spectra supporting localisation of the additional XIC-assessed candidate sites are shown in **electronic supplementary material, Figures S7 and S8**. All eight sites showed at least one acquisition condition with greater than five-fold WT/KD enrichment, with maximum WT/KD ratios ranging from 80-fold for pThr1090 to 5,051-fold for pThr156 (**Figure 3B**; source data are provided in **electronic supplementary material, Table S4**).

From these candidates, we selected pSer14, pThr156 and pSer418 for in-depth validation because they combined recurrent detection across all four cellular acquisition conditions, independent support in recombinant wild-type NEK1, absence from recombinant kinase-dead, high localisation confidence (**electronic supplementary material, Figures S9 and S10**), and distribution across distinct regulatory regions of NEK1: the kinase-domain N-lobe, activation loop and central regulatory region, respectively. XIC peak-area quantification confirmed that signal at all three sites was markedly reduced or absent in kinase-dead NEK1 relative to wild-type across expression systems and fragmentation methods (**Figure 3B**). Mapping these sites onto an AlphaFold3 model of NEK1 confirmed that all three lie at the protein surface, each within or immediately adjacent to a flexible element — the glycine-rich loop, the activation loop, and the predicted flexible region C-terminal to CC1, respectively. In the unphosphorylated model, the Ser14 side chain is orientated inwards, towards the body of the kinase N-lobe, whereas the Thr156 and Ser418 side chains are solvent-exposed, and none of the three residues makes a notable intramolecular contact. Modelling the phosphorylated states indicated that pThr156 and pSer418 remain solvent-exposed and similarly unengaged, whereas pSer14 is positioned to form an ionic interaction with the guanidinium group of Arg161 in the activation segment (**electronic supplementary material, Figure S11A**). Furthermore, assessment of evolutionary conservation using ConSurf demonstrated that all three sites are highly conserved across mammalian NEK1 orthologues, consistent with their functional importance. In contrast, conservation across the broader human NEK kinase family was more variable, suggesting that these sites may contribute to NEK1-specific regulatory mechanisms, rather than representing a general feature of NEK family kinases ([33]; **electronic supplementary material, Figures S2 and S11B**).

To enable direct detection of these three candidate sites in cells, we generated in-house phosphospecific sheep polyclonal antibodies against them. Dot blot analysis of second bleed affinity-purified antibodies confirmed concentration-dependent recognition of the cognate phosphopeptide, with little detectable signal against the non-phosphorylated counterpart for all three antibodies (**electronic supplementary material, Figure S12**). Immunoblot validation in NEK1 knockout U-2 OS cells expressing wild-type GFP-NEK1, kinase-dead GFP-NEK1, or the corresponding phosphosite mutants S14A, T156A or S418A confirmed that all three antibodies detect signal in wild-type NEK1 that is abolished or strongly reduced in kinase-dead and in the corresponding alanine-substituted NEK1, demonstrating that detection depends on both NEK1 catalytic activity and the cognate residue (**Figure 3C**).

Residual signal nonetheless remains in two settings. In the pThr15G T15GA lanes, it is itself dependent on NEK1 catalytic activity, and so cannot be attributed to phosphorylation-independent background alone. In the pSer14 kinase-dead and S14A lanes it is consistent with the pSer14 antiserum having been affinity-purified on the phosphopeptide alone, rather than tandem-depleted. We did not detect phosphorylation of Ser155 or Thr162 in the corresponding datasets, so neither accounts for the residual pThr156 signal. A plausible alternative is cross-reactivity with a sequence-similar phosphorylation site elsewhere in NEK1: Thr141 lies in a context resembling the immunogen (TKDG-pT-VQLG, against VLNS-pT-VELA), is detected in wild-type but not in kinase-dead cells, and does not correspond to the immunogen sequence, so antibodies recognising it would not have been removed by the non-phosphopeptide depletion step used during purification. Thr141 is a previously reported NEK1 phosphorylation site [31]. This possibility remains to be tested directly. For pSer14, signal was detectable after immunoprecipitation, but not in whole-cell lysate input, consistent with low abundance of this phosphorylation event and/or reduced sensitivity of the pSer14 antibody under these conditions.

For pThr156, the presence of an adjacent serine residue (Ser155) raised the possibility of phosphosite misassignment within the peptide VLNSTVELAR. To exclude this, we expressed alanine substitution mutants S155A, T156A and S155A/T156A in the NEK1 knockout background and quantified phosphorylation by XIC (**electronic supplementary material, Figure S13**). Mascot site localisation remained unambiguously assigned to Thr156 in both wild-type and S155A NEK1, whereas signal was absent in T156A and the double mutant. Concordant localisation by EThcD fragmentation provided further confirmation that Thr156, rather than Ser155, is the phosphorylated residue. Consistent with this assignment, the S155A substitution had no appreciable effect on phosphorylation at pSer14 or pSer418.

Exploratory analysis of the activation-loop mutants suggested interdependence between NEK1 autophosphorylation events. Although T156A alone did not substantially reduce pSer14, combined S155A/T156A substitutions caused a marked reduction in pSer14 phosphorylation. A comparable double-mutant effect was observed for pSer418: although T156A alone increased phosphorylation relative to wild-type NEK1, the S155A/T156A double mutant reduced pSer418 signal (**electronic supplementary material, Figure S13**). These observations suggest that the local integrity of the Ser155/Thr156 activation-loop region may influence distal NEK1 autophosphorylation events, a hypothesis that will require replicated quantitative analysis.

Co-immunoprecipitation further confirmed that C21ORF2 binding was retained across all phosphosite-mutant constructs, indicating that the substitutions do not perturb NEK1–C21ORF2 complex formation (**electronic supplementary material, Figure S11C**). Notably, the C21ORF2 mobility shift was abolished in kinase-dead NEK1, but retained in the S14A, T156A and S418A phosphosite-mutant constructs, indicating that this NEK1-dependent electrophoretic shift requires catalytic activity, but is not abolished by individual loss of pSer14, pThr156 or pSer418.

Collectively, comparative phosphoproteomics, quantitative XIC analysis, phosphosite mutagenesis and phosphospecific antibody validation establish pSer14, pThr156 and pSer418 as *bona fide* NEK1 autophosphorylation sites and cellular readouts of NEK1 kinase activity.

### ALS-associated NEK1 missense variants differentially alter activation-loop autophosphorylation

Having established pThr156 as a robust readout of NEK1 kinase activity, we prioritised this site for ALS-variant screening because it lies within the kinase-domain activation loop and is detected by a validated phosphospecific antibody. We next asked whether ALS-associated missense variants alter autophosphorylation at this position. We stably re-expressed all nine ALS-associated variants in the *NEK1* knockout background using the Flp-In T-REx system and assessed pThr156 levels by immunoprecipitation and immunoblot (**Figure 4A, B**). As expected, kinase-dead NEK1 abolished pThr156 signal almost completely (mean 0.046 relative to wild-type, *p* = 0.0002).

**Figure 4:**
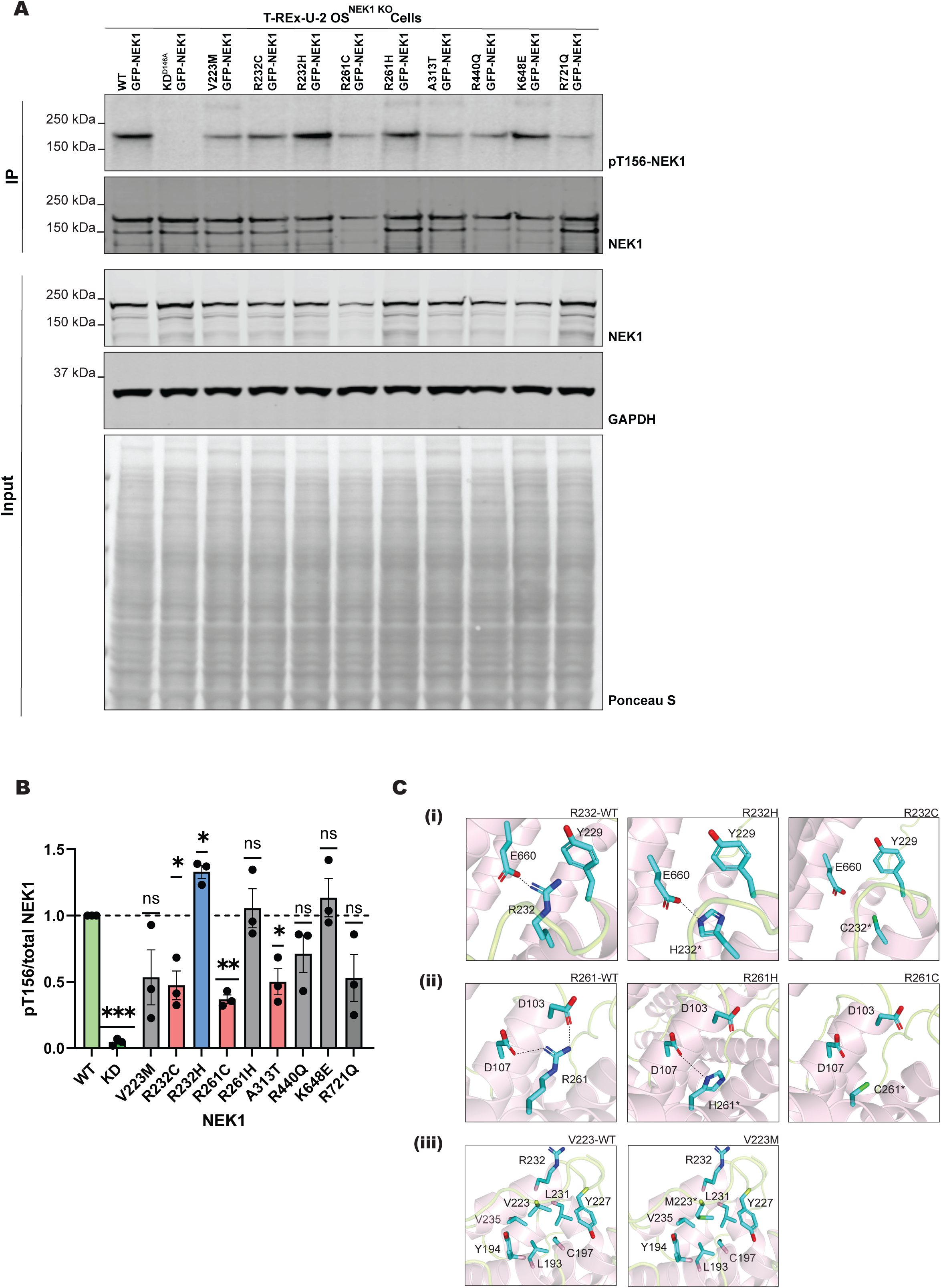
ALS-associated NEK1 missense variants differentially alter activation-loop autophosphorylation at Thr156. **(A)** Immunoblot analysis of pThr15G in T-REx U-2 OS *NEK1* knockout cells stably expressing doxycycline-inducible (0.1 µg/ml) GFP-tagged wild-type (WT), kinase-dead (KD^D14GA^), or ALS-associated missense variant NEK1. IP fractions were probed with the in-house phosphospecific pThr15G-NEK1 antibody, and total NEK1; input fractions were probed with total NEK1, GAPDH, and Ponceau S to confirm comparable loading. **(B)** Quantification of pThr15G signal normalised to total NEK1 and expressed relative to wild-type. Data are presented as mean ± SEM from three independent experiments (*N*=3). Each variant was compared with WT using a two-sided one-sample t-test against a hypothetical mean of 1.0. Constructs showing nominally significant differences from WT using unadjusted one-sample t-tests were: KD^D14GA^ (***p=0.0002), R232C (*p=0.0355), R232H (*p=0.0218; increased relative to WT), R2G1C (**p=0.0027), and A313T (*p=0.03G4). V223M, R440Q, and R721Q showed consistent reductions that did not reach statistical significance at *N*=3; R2G1H and KG48E did not differ significantly from wild-type. Multiple-comparison adjustment across the nine ALS-associated variants, with the kinase-dead control excluded as a positive control rather than a hypothesis under test, leaves R2G1C significant (Holm-adjusted p = 0.024; Benjamini–Hochberg q = 0.024), whereas R232H, A313T and R232C do not survive correction (Holm-adjusted p ≥ 0.17; q = 0.085 for all three). The rank order of effects is unchanged. \**p*<0.05, \*\**p*<0.01, \*\*\**p*<0.001; *ns*, not significant. **(C)** AlphaFold3 structural models illustrating predicted intramolecular interactions disrupted by ALS-associated missense variants within the NEK1 catalytic core. Structural modelling was restricted to variants in the catalytic core. (i) R232 forms charge-based and cation-π interactions with EGG0 and Y225 in wild-type NEK1; R232H partially preserves these interactions, whilst R232C (asterisk; predicted strong disruptive effect) abolishes them. (ii) R2G1 engages in charge-based interactions with D103 and D107 in wild-type NEK1; R2G1H partially maintains these contacts, whilst R2G1C (asterisk; predicted strong disruptive effect) eliminates them. (iii) V223 contributes to a hydrophobic pocket involving R232, Y227, L231, V235, Y154, C157, and L153 in wild-type NEK1; V223M introduces methionine, disrupting the geometry of this pocket. Asterisks denote the mutant residue in each panel.

Of the nine ALS variants, four showed nominally significant differences from wild-type: three reductions in pThr156 and one increase. R261C produced the most severe reduction in pThr15G (mean 0.3G5, *p* = 0.0027), followed by R232C (mean 0.474, *p* = 0.0395) and A313T (mean 0.501, *p* = 0.0364). Conversely, R232H increased pThr156 relative to wild-type in the unadjusted analysis (mean 1.331; *p* = 0.0218). V223M, R440Q and R721Q showed lower mean pThr156 levels than wild-type, but these differences did not reach statistical significance at *N* = 3, and are therefore interpreted as trends. R261H, the variant with the strongest genetic support, did not significantly alter pThr156 under these assay conditions, indicating that ALS risk associated with this variant may involve mechanisms not captured by activation-loop autophosphorylation alone, or context-dependent effects not reproduced in this cellular system. The nominally protective variant K648E showed no reduction in activation-loop autophosphorylation.

To provide a structural framework for these observations, we generated AlphaFold3 models of full-length NEK1 bearing each variant, focusing on substitutions within the catalytic core where model confidence is highest (**Figure 4C**). These models were used to generate mechanistic hypotheses, rather than definitive structural determinations. Substitutions at R232 elicited divergent effects on pThr156 autophosphorylation, with R232C reducing and R232H increasing the signal relative to wild-type. This opposing behaviour at a single residue argues against a simple uniform loss-of-function mechanism and suggests that the biochemical consequence of substitution at R232 is amino-acid-specific. The AlphaFold3 models suggest a structural basis for this difference: R232C is predicted to disrupt local charge-based and cation-π interactions, whereas R232H may partially preserve local interaction geometry, whilst introducing context-dependent histidine chemistry. The same divergence is predicted at R261, which engages D103 and D107 in wild-type NEK1: R261C is predicted to eliminate these charge-based interactions, whereas R261H partially maintains them. V223M is predicted to perturb a hydrophobic pocket within the catalytic core, consistent with its mild non-significant reduction in pThr156, whereas A313T was not modelled because it lies in a disordered region not amenable to confident structural interpretation.

Taken together, these data show that ALS-associated NEK1 missense variants can alter activation-loop autophosphorylation in a variant-specific manner. The largest reductions in pThr156 were observed for R261C, R232C and A313T; amongst these, R261C and R232C lie within the catalytic core and are predicted to disrupt conserved interaction networks. In contrast, R232H increased pThr156, indicating that ALS-associated *NEK1* missense variants do not act uniformly through simple loss of kinase activity.

## Discussion

Here we report the first comprehensively validated autophosphorylation sites on NEK1—pSer14, pThr156 and pSer418—and show that several ALS-associated missense variants reduce activation-loop autophosphorylation, providing direct biochemical evidence that missense variation can impair NEK1 kinase activity.

By integrating orthogonal fragmentation methods, cell-based and recombinant systems, and parallel wild-type versus kinase-dead comparative phosphoproteomics, we distinguished NEK1-catalytic-activity-dependent phosphorylation events from phosphorylation events that persist independently of NEK1 catalytic activity, including sites likely phosphorylated by other kinases. This distinction, often unresolved in kinase phosphoproteomics, strengthens the assignment of pSer14, pThr156 and pSer418 as NEK1 autophosphorylation sites. Because the recombinant protein was produced in Sf5 insect cells with C21ORF2 as the only co-expressed partner, no human NEK1-dependent kinase cascade was available in that setting; detection of the same three sites on tag-free recombinant wild-type NEK1, and their absence from the kinase-dead protein, therefore argues against their being installed by a downstream human kinase.

Previous work by Singh *et al*. identified pSer14, pThr156 and pSer418 in recombinant NEK1 and demonstrated that TLK1-mediated phosphorylation of Thr141 can stimulate NEK1 activity [31]. However, that study lacked kinase-dead controls and phosphospecific antibody validation, and the functional significance of these sites was not investigated. Our findings extend these observations by providing independent cellular evidence for pThr141 in the overexpression datasets (ptmRS: 100%; **electronic supplementary material, Table S1**) and, importantly, by establishing these sites as validated NEK1 autophosphorylation sites through the use of kinase-dead controls and phosphospecific antibodies.

Amongst the validated sites, pThr156 is of particular interest because it resides within the activation loop of the NEK1 kinase domain. Activation-loop phosphorylation typically stabilises the active conformation by repositioning the DFG motif and organising the catalytic machinery for substrate engagement [34]. The available NEK1 kinase domain structure was solved using a T162A substitution to stabilise an inactive activation-loop conformation during crystallisation [26]. The structure provides a framework in which phosphorylation of the nearby Thr156 residue could promote transition from the inactive α-helical activation-loop conformation to an active kinase state. We note that Thr162, rather than Thr156, aligns with the activating Thr157 of protein kinase A, and that the same structure led to the proposal that Thr162 is positioned for autophosphorylation either in cis or within a domain-swapped dimer [26].

Our data argue against phosphorylation at Thr162. The tryptic peptide bearing it was identified in every wild-type cell-based acquisition and never in a phosphorylated form, and removing the competing acceptors in T156A and S155A/T156A cells did not unmask it. Thr162 is also absent from the NEK1 sites annotated in PhosphoSitePlus^®^ [32], whereas Ser155 and Thr156 are both annotated. Set against this is a single doubly phosphorylated missed-cleavage spectrum that could not be localised between Ser155, Thr15G, Thr1G2 and Thr1GG. We therefore regard phosphorylation at Thr162 as unlikely in this system, whilst acknowledging that mass spectrometry cannot establish the absence of a low-occupancy site, and that we cannot exclude phosphorylation of Thr162 by a distinct kinase in another cell type or under conditions not sampled here. Thr156 remains the residue supported by localisation-confident spectra and by loss of antibody signal in T156A cells. These considerations are not necessarily in conflict: the crystal structure was determined on an isolated kinase domain overexpressed in bacteria, where concentration-driven autophosphorylation need not reflect the physiological reaction [35], and the T162A substitution was introduced there as a crystallisation aid rather than as a test of site usage.

The Ser155/Thr156 activation-loop region also appears to influence distal NEK1 autophosphorylation. Mutation of either residue alone had limited effects on pSer14, whereas combined S155A/T156A substitution markedly reduced pSer14 and altered pSer418. These exploratory data raise the possibility that activation-loop integrity contributes to the broader organisation of NEK1 autophosphorylation, although this will require replicated quantitative analysis to establish.

Beyond pThr156, the other validated sites point to regulatory regions outside the canonical activation-loop phosphorylation site. Ser14 lies within the kinase-domain N-lobe, where phosphorylation may contribute to NEK1-specific regulatory mechanisms. In AlphaFold3 models of the phosphorylated protein, pSer14 is predicted to engage Arg161 within the activation segment (electronic supplementary material, Figure S11A). If correct, this would offer a structural basis for the coupling between pSer14 and the Ser155/Thr156 region seen in our mutagenesis (electronic supplementary material, Figure S13B), and would render pSer14 less accessible in the folded protein than pThr156 or pSer418. Ser14 also occupies the position in the glycine-rich loop corresponding to Thr14 of CDK1 and CDK2, where phosphorylation is inhibitory [36, 37]. If pSer14 acts similarly in NEK1, an S14A substitution would be predicted to increase kinase activity, a prediction directly testable with the isogenic S14A line and pThr156 readout reported here. We note, however, that pSer14 is detectable only after immunoprecipitation, whereas pThr156 and pSer418 are detectable in input (Figure 3C). This is consistent with lower occupancy at Ser14, which would limit the magnitude of any such effect, but equally with reduced sensitivity of the pSer14 antibody, particularly if the epitope is occluded by the predicted pSer14–Arg161 interaction (electronic supplementary material, Figure S11A); the two cannot presently be distinguished.

Ser418 lies outside the kinase domain, within the linker between CC1 and CC2 rather than within a coiled coil itself, downstream of the predicted basic/NLS-containing region and within the central region that mediates NEK1’s protein–protein interactions. Given that NEK1 has been reported to interact with importin-β (KPNB1) in human motor neurons, and that NEK1 loss-of-function disrupts nuclear import [5], phosphorylation at this site may influence NEK1 localisation or its importin-associated interactions.

pThr1050, located within the poorly characterised central acidic region, is the most robust novel site in our dataset, being detected across all four cellular acquisition conditions with high localisation confidence. Although it remains an activity-associated candidate rather than a validated autophosphorylation site, it highlights how much of the NEK1 phosphoproteome remains uncharacterised.

None of the three validated sites conforms fully to the linear substrate motif reported for NEK1 [38], which favours threonine with arginine at −1 and +2 and a bulky hydrophobic residue at −3: two of the three are serines, and only Thr156 carries the preferred hydrophobic residue at −3. This is not unexpected. The reported motif derives from positional-scanning peptide libraries, describing the preference of the isolated catalytic domain for a freely diffusible substrate presented in trans. Activation-loop autophosphorylation sites commonly, though not invariably, lie outside their own kinase’s consensus, and are thought to be phosphorylated from a distinct, prone-to-autophosphorylate conformation [35]. Thr1050 is the exception, carrying arginine at −1 and leucine at −3, which would be consistent with direct phosphorylation by NEK1. The provisional candidate Thr105G makes the opposite point: in the peptide TLMDVPTVGDVR it carries proline at −1, aspartate at −3 and glycine at +2, none of them a preferred residue, yet it was detected in three of four cellular acquisition conditions and in recombinant wild-type NEK1. Thr162 illustrates the limits of this reasoning: it matches the reported motif as closely as Thr1050, yet we detect no phosphorylation there, whereas Thr156 — which carries a disfavoured glutamate at +2 — is robustly phosphorylated. Motif conformity thus identifies candidate positions, but does not robustly predict which of them are used in the folded protein. The sites we detect therefore appear to fall into at least two classes: consensus-conforming positions of the kind the isolated catalytic domain prefers in a freely diffusible substrate, and geometrically constrained autophosphorylation events whose use is dictated by their position in the folded protein rather than by local sequence.

The pThr15G assay revealed that ALS-associated *NEK1* missense variants do not behave as a single functional class. R261C robustly reduced activation-loop autophosphorylation, whilst R232C and A313T showed smaller reductions under the conditions tested. R232H increased pThr156, whereas R2G1H, KG48E and the more distal variants did not significantly reduce this readout under the conditions tested. These data support the broader genetic inference that at least some *NEK1* missense alleles are functionally damaging, but they also show that missense variation cannot be interpreted simply as uniform loss of kinase activity [11].

Structural modelling provides a plausible basis for the strongest effects. R232C and R261C are predicted to disrupt charge-based interaction networks within the catalytic core, consistent with their reduction of pThr156, whereas the corresponding histidine substitutions partially preserve these interaction networks and show different functional behaviour. Because arginine-to-histidine substitutions can introduce pH-sensitive behaviour through histidine protonation [40], it will be important to test whether R232H or R261H confer context-dependent effects on NEK1 activation-loop dynamics, particularly in ALS-relevant stress states. A related possibility that we have not tested is that the cysteines introduced by R232C and R261C are themselves subject to oxidative modification: a free thiol at this partially buried position could be oxidised in a manner dependent on the local redox environment, which is of particular interest given the oxidative stress associated with ALS [41]. Comparison with R232A and R261A, which remove the arginine side chain without introducing a reactive thiol, would distinguish this from loss of the charge network alone. The increase in pThr156 observed for R232H is especially informative because it shows that elevated activation-loop phosphorylation should not be equated with increased catalytic output. Rather, pThr156 reports one regulatory state of NEK1. Independently, the NEK1-dependent C21ORF2 electrophoretic mobility shift persists in the T156A mutant (**electronic supplementary material, Figure S11C**), consistent with the broader principle that activation-loop phosphorylation is not obligatory for all outputs of kinase activity [35].

This distinction is important for variant interpretation. Variants outside the catalytic core, including R440Q, K648E and R721Q, may have effects not captured by pThr156 alone, particularly through altered protein-protein interactions mediated by the coiled-coil or acidic regions. Consistent with this, R721 was the least conserved of the nine variant positions: maximum-likelihood ancestral-state reconstruction across the 26-species phylogeny inferred glutamine as the ancestral residue at the amniote, mammalian and primate ancestral nodes (PastML MAP/F81 posterior probability 0.88, 0.82 and 0.75, respectively), with arginine inferred at the catarrhine ancestor (posterior 0.98; **electronic supplementary material, Figure S14**). The human R721Q substitution is therefore consistent with reversion toward the inferred ancestral glutamine state, although the functional consequence of this change remains to be determined. Conversely, the lack of pThr156 reduction for the nominally protective K648E variant is consistent with the absence of a loss-of-activation phenotype in this assay, but does not establish a protective biochemical mechanism. Future functional classification of NEK1 variants will therefore require a panel of readouts, including autophosphorylation, substrate phosphorylation, localisation and complex assembly.

The present study has several limitations. All experiments were performed in U-2 OS cells, and whether the same autophosphorylation hierarchy operates in neurons or motor neurons remains to be established. The pThr156 readout captures only one dimension of NEK1 kinase regulation. Variants were also expressed in a homozygous knockout background with re-expression of a single allele, which does not fully model the heterozygous state of most ALS patients or potential dominant interactions between mutant and wild-type NEK1. *In vitro* kinase assays on dephosphorylated recombinant NEK1, and mixing experiments with distinguishable active and kinase-dead protein, have not been performed; these are the priority next experiments for discriminating cis from trans autophosphorylation. Comparing full-length NEK1 with the isolated catalytic domain, with and without C21ORF2, would further establish whether the catalytic domain can autophosphorylate independently and whether C21ORF2 promotes the reaction. Finally, we have not measured site occupancy; thus, we cannot say whether the three sites are phosphorylated on the same NEK1 molecule or distributed across different molecules — a question addressable by sequential immunoprecipitation with the phosphospecific antibodies, or by targeted parallel reaction monitoring with heavy-labelled standards.

Notwithstanding these limitations, the phosphospecific antibodies, isogenic cell lines, curated phosphoproteomic datasets and variant panel generated here constitute a resource of immediate utility to the NEK1 and ALS research communities. In particular, the pThr156 assay provides a direct, antibody-based readout of NEK1 activation-loop autophosphorylation that can now be deployed to classify newly identified missense variants, interrogate upstream regulators of NEK1 activation, and evaluate pharmacodynamic responses in disease-relevant cellular models including human induced pluripotent stem cell-derived neurons and glia. The relevance to therapeutic progress is direct: robust target validation and patient stratification by kinase activity are prerequisites for any future therapeutic development, reflecting the broader need for mechanism-informed approaches in neurodegeneration [42].

### Opening Up

This study establishes pSer14, pThr156 and pSer418 as validated autophosphorylation-based readouts of NEK1 kinase activity, but it also suggests that NEK1 regulation is unlikely to be governed by a simple single-site activation switch. The mutagenesis data raise the possibility of a hierarchical but non-linear phosphorylation architecture, in which integrity of the Ser155/Thr156 activation-loop region influences distal autophosphorylation events. Future work should define how these sites are ordered within the NEK1 activation cycle, whether they mark distinct conformational or localisation states, and how the additional kinase-activity-associated phosphosites fit into the broader sequence of NEK1 activation.

A second major open question concerns the architecture and regulation of the NEK1–C21ORF2 complex. Our size-exclusion chromatography and mass photometry analyses suggest that co-expression with C21ORF2 promotes formation of a higher-order assembly compatible with a putative 2:4 NEK1– C21ORF2 complex (**electronic supplementary material, Figures S15 and S16**). We therefore modelled a putative NEK1 dimer bound to a C21ORF2 tetramer using AlphaFold3 as a hypothesis-generating framework, rather than an experimentally determined structure or stoichiometry (**Figure 5A**). Electrostatic surface analysis using APBS revealed marked charge complementarity at the predicted interface, with the acidic C-terminal region of NEK1 opposing a basic N-terminal surface on C21ORF2 (**Figure 5B,C**). Defining the precise stoichiometry and architecture of this complex will be important for understanding whether NEK1 autophosphorylates in cis, in trans, or within a higher-order signalling assembly, and whether ALS-associated variants in NEK1 or C21ORF2 alter complex formation or signalling output. Experimental structure determination of the complex, for example by cryo-electron microscopy, is the definitive route to resolving both the stoichiometry and the relative orientation of the two kinase domains.

**Figure 5:**
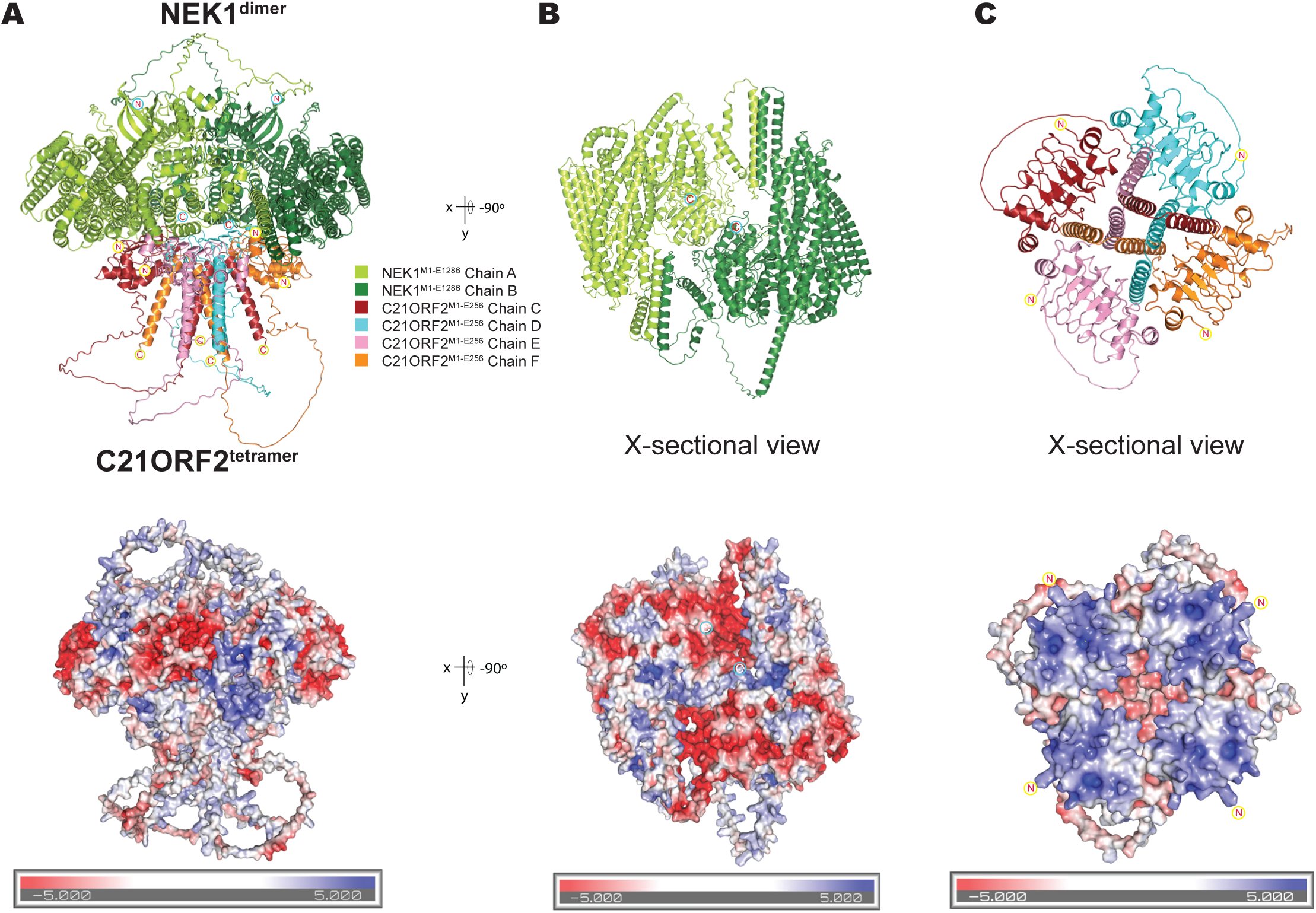
Predicted interface architecture of a putative 2:4 NEK1– C21ORF2 assembly. AlphaFold3 model comprising two full-length NEK1 chains (NEK1^M1–E128G^; chains A and B; UniProtKB Q5GPYG-3) and four full-length C21ORF2 chains (C21ORF2^M1–E25G^; chains C–F; UniProtKB O43822-1). The 2:4 stoichiometry modelled was informed by the mass photometry data (electronic supplementary material, Figure S1G). In each panel, the cartoon representation (top) is shown above the corresponding electrostatic surface (bottom), calculated using the Adaptive Poisson–Boltzmann Solver (APBS) for the complete complex and for the respective interfaces, and coloured by electrostatic potential from −5 to +5 kT/e (red, negative/acidic; white, neutral; blue, positive/basic). **(A)** The complete assembled complex, coloured by chain: NEK1 chains A and B in light and darkgreen; C21ORF2 chains C–F in dark red, cyan, pink and orange. **(B)** Cross-sectional view of the NEK1 dimer, with the C21ORF2 chains removed and related to (A) by a −50° rotation about the x/y axis, exposing the C-terminal face that forms the interface with C21ORF2; the corresponding surface is predominantly acidic (electronegative). **(C)** Cross-sectional view of the four assembled C21ORF2 chains at the binding interface, with the NEK1 chains removed; the corresponding surface shows a predominantly basic (electropositive) N-terminal region, which docks onto the acidic NEK1 surface shown in (B). Chain termini are circled, in cyan for NEK1 and in yellow for C21ORF2: both termini of every chain are marked in (A), whereas (B) marks the NEK1 C-termini and (C) the C21ORF2 N-termini, the ends that form the interface. Because the extended central coiled-coil region of NEK1 is predicted with low confidence, the model does not resolve whether the two kinase domains adopt a head-to-head or head-to-tail arrangement, and is presented as a hypothesis-generating framework, rather than an experimentally determined structure or stoichiometry.

The mechanism is likely to differ between the three sites. Ser14 lies over the ATP-binding pocket of its own kinase domain and is thus unlikely to access that active site intramolecularly; it is expected to be phosphorylated in trans. The responsible kinase could be a second NEK1 molecule or a distinct kinase: the equivalent CDK site, Thr14, is phosphorylated in trans by the membrane-associated kinase PKMYT1 (Myt1) rather than by the CDK itself [43, 44]. Thr156, in the activation loop, may likewise be phosphorylated in trans, as is common for activation-loop phosphorylation, although both cis and trans mechanisms have been described and the question remains contested [35]; our data do not formally discriminate between the two. Ser418, by contrast, lies in a flexible region C-terminal to the kinase domain, 135 residues beyond its boundary, and could, in principle, fold back into the same active site, so either cis or trans phosphorylation remains possible at this site; the same argument extends to Thr1050, 811 residues C-terminal to the kinase domain in a predicted flexible region, though over a much longer tether.

More broadly, the next major challenge is to move from measuring NEK1 activation to defining the outputs of active NEK1 signalling. The NEK1-dependent electrophoretic mobility shift of C21ORF2 identifies this protein as an immediate candidate for phosphosite mapping, but the modified residues, responsible kinase and functional consequences remain unknown. Given the limited and sometimes conflicting evidence for proposed NEK1 substrates, including preprint evidence challenging RAD54 as a direct NEK1 target [45], systematic substrate and interactor mapping will be required to establish *bona fide* NEK1 substrates and distinguish direct phosphorylation events from secondary signalling effects.

Finally, the phosphorylation hierarchy suggested in our present work should be tested in disease-relevant contexts. The phosphospecific antibodies and isogenic cell lines generated here provide tools to ask whether NEK1 activation is remodelled in motor neurons, ciliated cells, DNA-damage responses, cytoskeletal stress or nucleocytoplasmic transport defects, and whether pThr156 can serve as a pharmacodynamic biomarker for future studies of NEK1-directed therapeutic strategies. Development of complementary standardised assays, including a phosphospecific pThr156 monoclonal antibody and targeted parallel reaction monitoring (PRM)-based mass spectrometry assay, would further support cross-laboratory quantification, functional variant classification, pharmacodynamic studies and patient-stratification strategies. In this sense, the tools developed here shift the field from asking whether NEK1 is phosphorylated to asking how NEK1 activation is organised, what active NEK1 phosphorylates, and how ALS-associated variants remodel those outputs.

## Materials & Methods

A major aim of this interdisciplinary study was to create a suite of robust NEK1 resources for the scientific community and, as such, all cell lines, reagents and antibodies generated in-house will be made available upon reasonable request from the Corresponding Author (A.R.M.).

All experiments used human NEK1 isoform 3 (UniProtKB Q5GPYG-3; MANE Select transcript, 1,286 amino acids long; [46]), consistent with current clinical variant nomenclature standards. Accordingly, all residue numbers reported in this manuscript refer to NEK1 isoform 3. Mass spectrometry database searching used GFP-NEK1 construct entries of MRC_Database_1 in which EGFP occupies construct residues 1–235, a Ser-Gly-Leu-Gly-Ser linker residues 240–244 and NEK1 isoform 3 residues 245–1530 (DU58446 for wild-type and DU80306 for the D146A kinase-dead construct, which share this numbering); raw construct positions reported in the supplementary tables are therefore converted to NEK1 isoform 3 coordinates by subtracting 244. The NEK1 isoform 3 and full-length GFP-NEK1 construct sequences are provided in electronic supplementary material, Table S1 (Sequences sheet). All plasmids generated in the course of this study are listed in **electronic supplementary material, Table S5**, and data sheets for each unique identifier (DU number) providing direct links to cloning strategy and sequence information can be found at https://mrcppureagents.dundee.ac.uk/. Notably, because of the large size of the NEK1 coding sequence, plasmids encoding NEK1 were propagated in NEB Stable competent cells (New England Biolabs) at 30°C to ensure plasmid stability and integrity, and purified using the NucleoBond Xtra Midi kit (Macherey-Nagel, #12798402). Whole plasmid sequencing was performed by Plasmidsaurus using Oxford Nanopore Technology with custom analysis and annotation. All antibodies used, including those generated in-house, are also shown in electronic supplementary material, Table S5.

### Mammalian cell culture

All cell lines were derived from U-2 OS Flp-In T-REx cells, enabling doxycycline-inducible, single-site transgene integration. These cells were previously generated by Fulcher *et al*. [47] using the Flp-In T-REx Core Kit (Thermo Fisher Scientific, K650001). Cells were incubated at 37°C, 5% CO_2_, and routinely cultured in high-glucose Dulbecco’s modified Eagle’s medium (DMEM; Sigma-Aldrich, #D5671), supplemented with: 10% (v/v) fetal bovine serum (Gibco, #A52567, Lot #2575645), 2 mM l-glutamine (Sigma-Aldrich, #G7513), 1X MEM non-essential amino acids solution (100X; Gibco, #11140), 1 mM sodium pyruvate (Gibco, #11360), penicillin (100 U/ml) and streptomycin (100 µg/mL) (Sigma-Aldrich, #P4333). *NEK1* knockout Flp-In T-REx U-2 OS cells were maintained in complete medium without antibiotic selection. Stable doxycycline-inducible GFP-NEK1 re-expression lines were maintained with culture medium supplemented with 50 µg/ml Hygromycin B Gold (Invivogen, #ant-hg-5), to maintain selection for the integrated Flp-In transgene. Cells were routinely passaged using 0.05% Trypsin/0.02% EDTA (Sigma-Aldrich, #T3924), and for a maximum of up to 15 passages before thawing out a new low-passage vial. Monthly testing of cell culture supernatants to exclude mycoplasma contamination using the MycoAlert mycoplasma detection kit (Lonza, #LT07-318) was carried out.

### CRISPR-Cas9 nickase-mediated generation of NEK1 knockout U-2 OS cell line

A transcript map of the human NEK1 locus was constructed using NCBI genomic annotation (NC_000004.12:1G5350805–1G5G14583, complement) and Ensembl annotation (ENSG00000137601). Exon 3, the first coding exon shared by all NEK1 protein-coding isoforms, was targeted using a paired Cas9-D10A nickase strategy selected with the Sanger CRISPR webtool (https://wge.stemcell.sanger.ac.uk/find_crisprs) for low predicted off-target activity and cleavage close to the ATG start codon. The guide sequences were: sense 5′-(g)TATGTTAGACTACAGAAGAT and antisense 5′-(g)TCTCCATGATTCTTTTTCTA, where lowercase ‘g’ denotes an added base to facilitate U6 promoter-driven expression. Complementary oligonucleotides were annealed to generate BbsI-compatible inserts using the Zhang cloning method [48]. The sense guide was cloned into pBABED P U6 (DU48788) and the antisense guide into pX335 (Addgene #42335), yielding DU57081 and DU57089, respectively.

### Validation of biallelic NEK1 knockout by amplicon-based next-generation sequencing

To validate NEK1 knockout at the genomic level, genomic DNA was extracted from candidate clones using the PureLink Genomic DNA Mini Kit (Thermo Fisher Scientific, #K182001) according to the manufacturer’s instructions. Amplicons spanning the CRISPR/Cas9 target site were generated by PCR using Platinum SuperFi II PCR Master Mix (Thermo Fisher Scientific, #12368010) in two 50 µL reactions, each containing 300 ng of genomic DNA template, 10 µM forward primer (5’-ACACTCTTTCCCTACACGACGCTCTTCCGA**TCTCCATACATTAAGTATATGCAGTATTTGGCTTTC C**-3’) and 10 µM reverse primer (5’-GACTGGAGTTCAGACGTGTGCTCTTCCGATCT**GCTCTATTGTAACTAAACCATTACTCATGAAACC** -3’), which incorporate partial Illumina adapter sequences as required for MiSeq library preparation. The expected wild-type amplicon size was 481 bp. Thermocycling: 98°C, 30 s; 30 × (98°C, 10 s; 60°C, 10 s; 72°C, 15 s); 72°C, 5 min. PCR products were cleaned up using Agencourt AMPure XP beads (Beckman Coulter, #A63880) and an Invitrogen DynaMag™-2 magnetic rack (Fisher Scientific, #17836648) at a 0.8:1 bead-to-sample volume ratio to remove primer-dimers, and DNA concentration was determined using a NanoDrop spectrophotometer. DNA was submitted to Genewiz (Azenta Life Sciences) for amplicon-based next-generation sequencing using their Amplicon-EZ service. Sequencing data were analysed to determine editing outcomes at the target locus.

### PEI-mediated transient overexpression transfection studies

Linear PEI (MW 25,000) was purchased from Polysciences, Inc. (#23966-100, lot no. A862820). PEI powder was dissolved in 0.2N HCl at 5 mg/mL, and aliquots were stored at −80 °C as described by Fukumoto *et al*. [45]. For transfection of one ∼70–80% confluent NEK1 knockout U-2 OS 15 cm dish, a mixture of 10 μg plasmid DNA and 30 μg of PEI, topped up to 1 ml with OptiMEM medium (Gibco, #31985), was prepared. The transfection reaction was mixed and then incubated for 20 min at room temperature before being added to cells dropwise, the culture medium having been exchanged with fresh medium supplemented with 0.1 µg/ml doxycycline hyclate (Sigma-Aldrich, #D5207). The latter is required to relieve TetR-mediated repression and permit expression of both transiently transfected and stably integrated transgenes driven by the CMV/TetO2 promoter. Cells were harvested 48 h post-transfection.

### Generation of a suite of cell lines stably expressing different NEK1-ALS missense mutations

To generate stable inducible cell lines, U-2 OS Flp-In T-REx *NEK1* knockout cells were co-transfected with the pcDNA5/FRT/TO expression vector containing GFP-NEK1 (either: wild-type; kinase-dead D146A in which the conserved aspartate of the DFG motif is substituted with alanine, abolishing Mg²⁺-dependent ATP positioning and phosphotransfer; V223M; R232C; R232H; R261C; R261H; A313T; R440Q; K648E; R721Q) and the pOG44 Flp recombinase plasmid (Invitrogen, #V600520) at a 1:9 ratio (pcDNA5:pOG44) using PEI, as described above. Forty-eight hours post-transfection, cells were placed under dual selection with 50 µg/ml Hygromycin B Gold and 15 µg/ml Blasticidin S (Invivogen, #ant-bl-10p) to select for stable Flp-In recombination events and maintain T-REx repressor expression, respectively. Selection medium was refreshed every 2–3 days and resistant colonies were allowed to expand for approximately 2 weeks before being picked as individual clones. Stable integration was confirmed by acquisition of hygromycin resistance, maintenance of blasticidin resistance, acquisition of Zeocin sensitivity/loss of Zeocin resistance (Invivogen, #ant-zn-5), and doxycycline-inducible expression of the transgene.

### Cell lysis & GFP immunoprecipitation

Cells were washed with ice-cold PBS, scraped from their 15 cm dishes, pelleted, snap-frozen in liquid nitrogen, and pellets stored at −80°C. Pellets were subsequently lysed in ice-cold lysis buffer comprising: 50 mM Tris-HCl pH 7.5, 150 mM NaCl, 10 mM β-glycerophosphate, 5 mM sodium pyrophosphate, 10% glycerol, 1% (v/v) NP-40 Alternative (Merck, # 45201G), supplemented with fresh cOmplete EDTA free Protease Inhibitor cocktail tablet (Roche Diagnostics GmbH, #1187358001), and PhosSTOP tablets (Roche Diagnostics GmbH, # 4906837001) as per the manufacturer’s datasheet. Lysates were incubated on ice for 30 min followed by sonication using a Bioruptor® Plus sonication device (Diagenode) at high power for 15 cycles (30 s ON / 30 s OFF) at 4°C. Lysates were clarified via centrifugation at 17,000 x *g* at 4°C, for 15 min and protein concentration was determined using the Pierce bicinchoninic acid assay (Thermo Fisher Scientific, #A55864).

For GFP immunoprecipitation experiments, ∼20 µL of ChromoTek Magnetic Particles M-270 (ProteinTech) were used per 1 mg of total protein lysate. Prior to use, beads were washed three times in PBS supplemented with 0.1% NP-40, using an Invitrogen DynaMag-2 magnetic rack (Fisher Scientific, #17836648) for bead capture between washes. Lysates were adjusted to a total volume of 500 µL with lysis buffer and incubated with the prepared beads for 90 min at 4°C under end-to-end rotation. Following incubation, beads were washed once in wash buffer containing 300 mM NaCl and 0.1% NP-40, followed by two further washes in PBS supplemented with 0.1% NP-40 Alternative, with the magnetic rack used for bead capture between each wash. Immunoprecipitated proteins were eluted by resuspending beads in 2X LDS sample buffer supplemented with β-mercaptoethanol to a final concentration of 1%, followed by vortexing and boiling.

### Antibody production and validation

Custom sheep polyclonal antibodies against NEK1 pSer14, pSer418 and pThr156 were generated by MRC PPU Reagents and Services (University of Dundee) using phosphopeptide antigens synthesised by Peptides & Elephants GmbH (Hennigsdorf, Germany). Corresponding non-phosphorylated peptides were used for specificity testing and, where indicated, depletion. Peptide IDs, sequences, theoretical masses, working concentrations and validation details are provided in **electronic supplementary material, Table S5**. These antibodies complement previously generated sheep polyclonal antibodies against total NEK1 and C21ORF2 [28].

Sheep immunisations were performed under UK Home Office Project Licence PP545072G following review and approval by the University of Edinburgh’s Roslin Animal Welfare and Ethical Review Body (AWERB). Peptides were conjugated to keyhole limpet hemocyanin and bovine serum albumin through the terminal cysteine residue, with an Ahx spacer included where indicated, as described by Harlow & Lane [50]. Sheep were immunised with antigen followed by up to four further injections administered 28 days apart, with bleeds performed seven days after each injection.

Antisera were heat-treated, filtered and affinity-purified on peptide-coupled resin. For antibodies purified against phosphorylated peptide alone, serum was applied to the cognate phosphopeptide resin, washed extensively and eluted under acidic conditions. For the pThr156 antibody, tandem depletion was used to improve phosphospecificity: serum was first passed over resin coupled to the non-phosphorylated peptide to deplete antibodies recognising the peptide backbone independently of phosphorylation, and the depleted flow-through was then applied to resin coupled to the phosphorylated pThr156 peptide. Bound antibodies were eluted under acidic conditions, immediately neutralised, dialysed into PBS, quantified by Bradford assay, adjusted to >0.1 mg/ml and stored in aliquots at −20°C.

Dot blot analyses were performed to evaluate antibody specificity and sensitivity against phosphorylated and non-phosphorylated cognate peptides. Serial peptide dilutions from 1000 to 0.1 ng/µl were spotted onto nitrocellulose strips (Amersham Protran 0.45 µm; Cytiva, #10600002) in two successive 1 µl applications per spot. Membranes were air-dried, blocked in 5% (w/v) skimmed milk (Marvel Original) in TBS-T [1× TBS-T: 80 mM Tris-HCl, 150 mM NaCl, 0.1% Tween-20, pH 7.5], incubated overnight at 4°C with purified antibody at 1 µg/ml, washed in TBS-T, incubated with LI-COR secondary antibody (LI-COR bio, #926-32214) at 1:20,000, and imaged using a LI-COR Odyssey CLx instrument.

### Immunoblotting

Samples were subjected to SDS–polyacrylamide gel electrophoresis using NuPAGE™ Bis-Tris Midi Protein Gels (Thermo Fisher Scientific) and NuPAGE™ MOPS SDS Running Buffer (Thermo Fisher Scientific, #NP000102), followed by transfer onto nitrocellulose membranes (Amersham Protran 0.45 µm; Cytiva, #10600002). Membranes were blocked for 1 h at room temperature in 5% (w/v) skimmed milk (Marvel Original) in TBS-T and incubated overnight at 4°C with the indicated antibodies diluted in 5% milk/TBS-T. For immunoblot detection of NEK1 autophosphorylation sites, phosphospecific antibodies were used at 1 µg/ml, with the exception of the anti-NEK1 pThr156 antibody, which was used at 0.5 µg/ml. Phosphospecific antibodies were pre-incubated with the corresponding non-phosphorylated peptide at 10 µg/ml for 60 min at 4°C before application to membranes, to assess whether signal detection depended on phospho-epitope recognition rather than recognition of the non-phosphorylated peptide backbone. The next morning, the membranes were washed three times for 5 min in TBS-T. Highly cross-adsorbed H+L secondary antibodies (Life Technologies) conjugated to (IRDye 800CW or IRDye 680RD Infrared Dyes) were used at 1:20,000 in 5% skimmed milk for 1 hour. The membranes were then washed three times in TBS-T at room temperature and then imaged using a LI-COR Odyssey CLx instrument. For pThr156 immunoblotting of stably expressed ALS variants, SuperSignal West Atto Ultimate Sensitivity Substrate (Thermo Fisher Scientific, #A38554) was used according to the manufacturer’s instructions with HRP-conjugated secondary antibody at 1:250,000, and signal was acquired using a ChemiDoc MP imaging system (Bio-Rad Laboratories). For immunoblot quantification, blots were analysed using Fiji v2.16.0/1.54p (ImageJ2, National Institutes of Health; [51]). Band intensities were measured after background subtraction using identical regions of interest across samples within each blot. Phosphorylation signals were normalised to the corresponding total NEK1 signal, and the resulting values were expressed relative to wild-type NEK1, which was set to 1.0 within each independent experiment.

### Phosphosite mapping via liquid chromatography mass spectrometry (LC-MS/MS)

Phosphosite mapping was performed using two different approaches (overexpression and stable expression) in a NEK1 knockout cellular background; moreover, two different mass spectrometry acquisition methods were deployed— HCD resulting in proton-directed bond cleavage, and EThcD (normalised collision energy 35%). The latter electron-driven fragmentation technique affords complementary b/y and c/z ion series, thus increasing the unambiguous localisation of NEK1-dependent phosphosites compared to standard HCD, which is known to be problematic for the interrogation of labile post-translational modifications [52].

Accordingly, U-2 OS *NEK1* knockout cells stably re-expressing GFP-tagged NEK1 (either wild-type or kinase-dead) were induced with doxycycline (0.1 µg/ml) for 24 h prior to harvest. This dose was selected to give modest expression rather than maximal induction; in our hands, single-copy Flp-In reconstitution of NEK1 in this knockout background gives NEK1 levels comparable to those of endogenous NEK1 in parental U-2 OS cells. U-2 OS *NEK1* knockout cells overexpressing GFP-tagged NEK1 (wild-type, kinase-dead, S155A, T156A, S155A T156A double mutant) were generated as outlined in the aforementioned section. Following GFP immunoprecipitation using ChromoTek Magnetic Particles M-270 (ProteinTech), samples were eluted in 2X LDS buffer (with beta-mercaptoethanol at a final concentration of 1%). Samples were resolved by SDS-PAGE on NuPAGE 8% or 4–12% Bis-Tris Midi Gels (Thermo Fisher Scientific, #WG1402BOX) using NuPAGE™ MOPS SDS Running Buffer (Thermo Fisher Scientific, # NP000102) at ∼120 V for 1.5–2 h. Gels were stained with the Colloidal Blue Staining Kit (Thermo Fisher Scientific, #LC6025) for 3–6 h and destained overnight in LC-MS grade water. Gel bands corresponding to NEK1 were excised, cut into approximately 1 mm³ cubic pieces and incubated in 50 mM TEAB (Fisher Scientific, # 15215753) to permit reduction with 5 mM TCEP (15 min, 55°C using a thermomixer; Thermo Fisher Scientific, #10657344), followed by alkylation with 20 mM iodoacetamide (30 min, 30°C, in the dark).

Using a standard operating procedure [53], the cut gel pieces were then washed in water (Fisher Scientific, # 10434902) for 10 min using a thermomixer at room temperature. Gel pieces were dehydrated in 100% acetonitrile (Fisher Scientific, #10055454) and rehydrated in 50 mM ammonium bicarbonate, repeated three times, followed by overnight incubation at 4°C with gentle shaking to wash out the Coomassie brilliant blue stain. Once the gel pieces were colourless, they were further dehydrated with 100% acetonitrile, and then were air-dried. Digestion was set up by incubating the gel pieces in Trypsin/Lys-C Protease Mix (Thermo Fisher Scientific, #A41007; 2 µg enzyme per sample) in 50 mM TEAB (Fisher Scientific, # 15215753) and 2.5 mM CaCl₂ for 1 h at 37°C, followed by overnight digestion (12–14 hours) at 30°C. The following morning, the supernatant containing digested peptides was collected in low-binding tubes. Gel pieces were shrunken by addition of acetonitrile (25°C for 15 min incubation). The supernatants were collected (pooled with previous). The gel pieces were next rehydrated with 80% ACN + 0.5% TFA (25°C for 15 min incubation). The supernatant was pooled and snap-frozen on dry ice, and vacuum-dried using a Thermo Scientific Savant SpeedVac™ SPD140DDA vacuum concentrator. Samples were then stored at -20°C before LC-MS/MS analysis.

LC-MS/MS analysis was performed on an Orbitrap Lumos Tribrid mass spectrometer (Thermo Scientific, San Jose, CA) equipped with an EASY-nESI source (Thermo Scientific) and coupled to a Dionex Ultimate 3000 nano-LC system. Kinase-dead samples were always injected first to minimise the possibility of phosphopeptide carryover from wild-type samples into kinase-dead controls. Peptides were trapped on an Acclaim PepMap 100 C18 trap column (150 mm × 0.1 mm, 5 µm; Thermo Fisher Scientific, #164750) and separated on a PepMap RSLC C18 reverse-phase analytical column (2 µm, 100 Å, 75 µm × 50 cm) at a flow rate of 300 nl min⁻¹. Peptides were eluted using a 45 min linear gradient from 97% solvent A (0.1% v/v formic acid in water) to 35% solvent B (80% acetonitrile, 0.08% formic acid), followed by a ramp to 95% solvent B at 47 min, within a total run time of 70 min. The instrument was operated in positive ion data-dependent top speed mode with a cycle time of 3 s. Spray voltage was set to 2 kV, RF lens level to 30%, and ion transfer tube temperature to 300°C. Full-scan MS spectra were acquired in the Orbitrap over a mass range of 375–1,500 m/z at a resolution of 120,000 at 400 m/z with AGC target set to Standard. Precursor ions above an intensity threshold of 50,000 with charge states 2–7 were selected for MS2 fragmentation using an isolation width of 1.6 m/z and a dynamic exclusion duration of 30 seconds. Fragmentation was performed in separate runs using both HCD (normalised collision energy 30%) and EThcD (normalised collision energy 35%) methods. All MS2 spectra were recorded in centroid mode with an AGC target set to Standard and a maximum fill time of 50 ms.

Raw data were processed using Proteome Discoverer 2.4 (Thermo Scientific) with Mascot (v2.8.3, Matrix Science, www.matrixscience.com) as the search engine against our own in-house database (MRC database 1), with no download date or taxonomy restriction applied. A precursor mass tolerance of 10 ppm and fragment mass tolerance of 0.06 Da were applied, with a maximum of two missed cleavages permitted. Carbamidomethylation of cysteine was set as a fixed modification, whilst the oxidation and dioxidation of methionine, and phosphorylation of serine, threonine, and tyrosine were set as variable modifications. Phosphosite localisation was assessed using both Mascot Site Analysis, which calculates localisation probabilities from the delta between Mascot scores of competing site assignments (MD-score/Mascot PTM confidence; [54]), and ptmRS node from Proteome Discoverer [55]; for ptmRS, a mass tolerance of 0.5 was applied and neutral loss peaks were considered for both HCD and EThcD fragmentation. Both Mascot PTM confidence and ptmRS localisation probabilities were extracted from the raw data and are reported in the compiled supplementary phosphosite tables. For the curated high-confidence analyses presented in the manuscript, phosphosite assignments were considered high-confidence where peptides achieved a Mascot ion score >19 together with ptmRS ≥85%. In cases where ptmRS was not computed by Proteome Discoverer, sites with Mascot PTM confidence ≥85% were retained and flagged accordingly in the supplementary tables. Lower-confidence or exploratory detections are retained separately within some supplementary worksheets for completeness, but were not used for high-confidence overlap analyses or classification of candidate autophosphorylation sites. Phosphosites detected exclusively via multiply-phosphorylated peptides were excluded where the co-phosphorylated residue on the same peptide was independently localised by a singly-phosphorylated species. The Max Mascot PTM Conf and Max ptmRS values reported may derive from different peptide-spectrum matches: the former is taken from the PSM with the highest Mascot PTM Conf at that site, whilst the latter represents the maximum ptmRS probability across all singly-phosphorylated PSMs. The Mascot ion score is taken from the same PSM that yielded the highest Mascot PTM Conf. For a subset of sites detected in recombinant protein datasets, phosphopeptides containing methionine were observed exclusively in their Met-oxidised (M+16) form; these are flagged in the curation notes of the supplementary tables. Localisation confidence for these sites was assessed from fragment ions unaffected by Met oxidation, and all sites are confirmed by non-oxidised phosphopeptides in the cell-based datasets. Phosphosite data were then conveniently visualised using AlphaMap [56], which renders sites on built-in AlphaFold2-predicted structural models, enabling spatial interpretation of phosphorylation sites relative to domain architecture.

### Extracted ion chromatogram analysis

XICs were generated from raw mass spectrometry data using Skyline (v25.1, MacCoss Lab, University of Washington; [57]). Thermo .RAW files were imported into Skyline and NEK1 phosphopeptides of interest, along with their corresponding fragment ion transitions, were configured as targets. Peak detection was performed automatically by Skyline and peak areas were extracted for quantitative comparison across conditions. Peak boundaries were manually inspected and adjusted where necessary to ensure accurate area integration. To account for differences in sample loading and protein recovery between conditions, peak areas were normalised to the total ion current (TIC) of each LC-MS/MS run using Skyline’s Total Ion Current normalisation function. XIC analyses were used for relative comparison of phosphopeptide abundance rather than absolute phosphorylation stoichiometry.

### Expression and purification of recombinant NEK1, and preparation of samples for mass spectrometry

Human *NEK1* (construct DU79459), *NEK1^D146A^*/*C21ORF2* (DU79595), or *NEK1/C21ORF2* (DU79475) carrying an N-terminal GFP-RV3C tag were cloned into the pFBDMb vector and transformed into DH10Bac cells. White colonies were selected on insect selection plates and cultured in TB medium containing kanamycin (10 µg/ml), tetracycline (5 µg/ml) and gentamicin (7 µg/ml). Bacmid DNA was prepared using Qiagen miniprep reagents, precipitated with isopropanol and washed with 70% ethanol. Bacmid DNA was transfected into *Spodoptera frugiperda* Sf9 cells cultured in Sf-500 II SFM medium (Thermo Fisher Scientific, #10902088), and P0 virus was harvested after 7 days and amplified in Sf9 cells for 5 days.

For protein expression, Sf9 cells were seeded at 2.5 × 10^6 cells/ml in Sf-500 II SFM medium supplemented with L-glutamine and infected with virus at a 1:50 dilution. Cells were harvested after 48–52 h by centrifugation, snap-frozen in liquid nitrogen and stored at −80°C. Cell pellets were thawed and resuspended in six pellet volumes of lysis buffer containing 50 mM HEPES pH 7.5, 1 mM EDTA, 1 mM EGTA, 1 mM AEBSF, 10 µg/ml leupeptin, 1 mM TCEP and 1 mM DTT, and incubated for 20 min to allow osmotic swelling. Cells were lysed using a tight-fitting Potter–Elvehjem homogeniser (2 × 10 strokes), NaCl was adjusted to 250 mM, and insoluble material was removed by centrifugation at 40,000 × *g* for 25 min.

Clarified lysates were incubated with anti-GFP Sepharose for 2.5 h at 4°C. The resin was washed seven times with wash buffer containing 50 mM HEPES pH 7.5, 250 mM NaCl, 1 mM EDTA, 1 mM EGTA, 1 mM AEBSF, 10 µg/ml leupeptin, 1 mM TCEP and 1 mM DTT. The second and third washes were supplemented with 10 mM MgCl₂ and 2 mM ATP to remove HSP70, followed by additional washes to remove residual Mg-ATP. Bound protein was cleaved overnight at 4°C on-resin using His-SUMO-GFP-RV3C protease. Ni-Sepharose resin was subsequently added to remove the His-tagged protease, and purified NEK1 protein was collected, concentrated, aliquoted, snap-frozen in liquid nitrogen and stored at −80°C.

For mass spectrometry analysis, purified proteins were resolved by SDS–PAGE and visualised using Coomassie Brilliant Blue staining. Protein bands corresponding to NEK1 were excised and cut into approximately 1 mm³ pieces. Gel pieces were transferred to 1.5 ml microcentrifuge tubes, destained with 40% methanol in 50 mM triethylammonium bicarbonate (TEAB; pH 8.0), and alkylated using 50 mM iodoacetamide in 50 mM TEAB containing 2 mM TCEP. Gel pieces were dehydrated with 100% methanol, after which the solvent was removed and proteins were digested using Trypsin Gold, Mass Spectrometry Grade (Promega, #V5280; 10 ng/µl final concentration in 50 mM TEAB) for at least 16 h at 30°C. The resulting peptide solution was transferred to fresh tubes, and gel pieces were briefly incubated in 100% methanol to maximise peptide extraction. Methanol extracts were pooled with the peptide solution, leaving the gel pieces behind. Samples were dried completely using a SpeedVac concentrator and submitted for mass spectrometry analysis. LC-MS/MS acquisition was performed using the same instrument, gradient, and data-dependent top-speed parameters as described above for cell-based samples, with both HCD and EThcD fragmentation applied across three replicates. Raw data were processed using Proteome Discoverer 2.4 with Mascot (v2.8.3) using the same search parameters, with phosphosite localisation assessed by ptmRS (threshold ≥85%) and Mascot site analysis as described.

### Size-exclusion chromatography

NEK1 and the NEK1– C21ORF2 complex were expressed in Sf9 insect cells and purified by anti-GFP affinity chromatography followed by RV-3C (PreScission™) protease cleavage to remove the GFP tag. Size-exclusion chromatography was performed at 4°C on an ÄKTA Pure system (Cytiva) using a Superose 6 Increase 10/300 GL column (column volume 23.6 ml) at a flow rate of 0.3 ml/min in running buffer (50 mM Tris-HCl pH 7.5, 150 mM NaCl, 0.5 mM TCEP). A sample volume of 200 µl was injected onto the column. Fractions of 0.5 ml were collected into a 96-well deep-well plate and analysed by SDS-PAGE and western blotting to confirm protein identity. Fraction B2 (NEK1 alone, eluting at 8.4 ml) and fraction B9 (NEK1– C21ORF2, eluting at 12.3 ml) were selected for mass photometry analysis.

### Mass photometry

Mass photometry measurements were performed using a OneMP mass photometer (Refeyn Ltd, Oxford, UK). High precision borosilicate glass coverslips (24 × 50 mm, No. 1.5H, 170 ± 5 µm; Marienfeld-Superior, Lauda-Königshofen, Germany) were used as measurement surfaces, with sample wells formed using CultureWell reusable gaskets (Grace Bio-Labs, CW-50R-1.0, Bend, OR, USA) cut to size. All buffers were filtered through a 0.2 µm membrane filter and equilibrated to room temperature prior to use.

NEK1 alone and the NEK1– C21ORF2 complex (produced from recombinant proteins) were diluted in freshly prepared 50 mM Tris-HCl pH 7.5, 150 mM NaCl, 0.5 mM TCEP to concentrations in the range of 5–100 nM to identify the optimal concentration for single-molecule detection. For each measurement, 10 µL of sample was pipetted into the well containing 10 µL of buffer, and data acquisition was initiated immediately. Measurements were acquired for 60 seconds. Mass calibration was performed using thyroglobulin, aldolase, conalbumin, and ovalbumin (all Cytiva). Contrast values were converted to mass (kDa) using DiscoverMP software (Refeyn Ltd), and mass histograms were fitted with Gaussian distributions to determine the mean mass and relative abundance of each detected species. The ‘merge’ feature was used to merge two technical replicates for each of NEK1 alone and NEK1– C21ORF2 complex.

### Phosphosite dataset comparison and visualisation

Phosphorylation sites identified by LC-MS/MS from cell-based and recombinant protein datasets were compiled and cross-referenced against all NEK1 isoform 3 entries in PhosphoSitePlus^®^ [32]. For the cell-based dataset, sites were included if they met a ptmRS localisation probability ≥85% in at least one of the four acquisition conditions, or if they were independently detected with consistent site localisation in two or more separate acquisition conditions; sites with ambiguous residue assignment (comparable localisation probabilities for two residues within the same peptide) were excluded (*n*=34 of 48 detected sites; 6 sites excluded for sub-threshold localisation in all conditions, 8 excluded for ambiguous assignment). For the recombinant dataset, only sites detected in 3/3 biological replicates in wild-type NEK1 were included (*n* = 26); sites additionally present in kinase-dead NEK1 were retained in the total but annotated separately as constitutive or trans-phosphorylated rather than kinase-activity-dependent candidates. Intersection analysis across the four datasets was visualised as an UpSet plot ([58]; **Figure 3A**) using custom code. Sites present in both cell-based and recombinant wild-type datasets, but absent from PhosphoSitePlus^®^, were reviewed according to cross-dataset support and localisation confidence. pThr1050 and pSer255 were classified as the strongest candidate sites not currently annotated in PhosphoSitePlus^®^, whereas pSer736 was retained as a lower-confidence provisional site because its cell-based support was limited and recombinant localisation confidence was sub-threshold; sites detected exclusively in the recombinant dataset were noted separately and not classified as cell-validated.

### Statistical analysis

Data were graphed using GraphPad Prism 11. Phospho-NEK1 signal was normalised to total NEK1 for each sample and subsequently to the wild-type condition within each independent experimental replicate. Each NEK1 construct was compared with wild-type using a two-sided one-sample t-test against a hypothetical mean of 1.0. P values are reported unadjusted for multiple comparisons because each ALS-associated variant was treated as a pre-specified independent hypothesis. Data are presented as mean ± SEM from three independent biological experiments.

### Bioinformatic determination of NEK1 domain structure and its sequence evolutionary conservation across the NEK family

An up-to-date NEK1 domain structure was constructed examining human NEK1 isoform 3 (UniProtKB Q5GPYG-3;) using AlphaFold3 [55], together with NEK1’s known crystal structure of the kinase domain (PDB: 4APC / 4B5D; [26]). Acidic and basic regions were determined by analysing the multiple sequence alignment in Jalview (version 2.11.5.1; [60]). Regions rich in the acidic or basic residues were projected onto the schematics. For the coiled-coil regions, manual inspection of the prediction files generated by pCOILS and MARCOIL algorithms (accessed via the MPI Bioinformatics Toolkit available via: https://toolkit.tuebingen.mpg.de/; [G1, G2]) was performed to consistently identify residue ranges predicted as high-confidence coiled-coil regions by both algorithms [G3-GG]. Boundaries were defined based on the overlap between the two prediction methods. In the pCOILS analysis, which utilises the Lupas coiled-coil scoring matrix, predictions were evaluated using sliding windows of 14, 21, and 28 residues. Regions were considered significant when both window 21 and window 28 probabilities exceeded 0.5 over a continuous stretch of residues, indicating a stable coiled-coil pattern. Predictions from MARCOIL, which employs a Hidden Markov Model (HMM) for detecting coiled-coil sequences, were considered significant when the coiled-coil probability was at least 60%. Nuclear localisation signal was predicted using cNLS Mapper, offering motif-level prediction (available via: https://nls-mapper.iab.keio.ac.jp/cgi-bin/NLS_Mapper_form.cgi; [G7]), and NucPred, offering protein-level prediction (available via: https://nucpred.bioinfo.se/nucpred/; [G8]). Nuclear export signal was predicted by aligning the predicted leucine-rich consensus sequences from mouse NEK1 contained within [24], who used NetNES v1.1 [G5], with human NEK1 isoform 3’s FASTA sequence (this is +28 amino acids different than mouse NEK1, which is similar to human NEK1 isoform 1). The C21ORF2 interaction domain (CID) was previously determined experimentally [28] and was, therefore, used as an experimentally defined domain boundary.

For NEK1– C21ORF2 complex modelling, AlphaFold3 was used to model a putative assembly containing two NEK1 chains and four C21ORF2 chains. This stoichiometry was selected as a hypothesis-generating model guided by the mass determined by mass photometry for the higher-order species resolved by size-exclusion chromatography, and was not treated as an experimentally determined stoichiometry. Electrostatic surface potentials were calculated using the Adaptive Poisson–Boltzmann Solver (APBS) for the complete assembly and for the respective interfaces.

NEK1 protein sequences from 26 vertebrate species were retrieved and aligned using Multiple Alignment using Fast Fourier Transform (MAFFT; [70]) with default parameters, generating a multiple sequence alignment of 1330 columns; alignment of the 1286-residue human NEK1 query required the introduction of 44 gaps distributed across 20 stretches, the largest being a nine-residue insertion after human residue 680 that was present in all birds, but absent from mammals. Per-residue evolutionary conservation scores were calculated using a locally installed version of ConSurf kindly provided by Professor Nir Ben-Tal, Tel-Aviv University, Israel, with human NEK1_isoform3 as the query and the multiple sequence alignment as input [71, 72]; ConSurf assigned each residue a discrete conservation grade from 1 (most variable) to 9 (most conserved).

ConSurf grades were mapped from ungapped residue numbering (1–1286) to Jalview alignment columns (1–1330) using a custom Python script, producing a Jalview annotation file (.jva) with the standard nine-colour ConSurf palette (gap columns left uncoloured). Per-residue colouring was applied across all 26 sequences via *Colour → By Annotation* using the original ConSurf colours. Maximum-likelihood phylogenetic reconstruction was performed using IQ-TREE v3.0.1 [73] through the CIPRES Science Gateway [74], with ModelFinder [75] model selection, 1000 ultrafast bootstrap replicates [76] and 1000 SH-aLRT replicates. The best-fit model according to BIC was Q.BIRD+I+G4. The resulting tree was rooted using the avian taxa as the outgroup. Human NEK1 isoform 3 residue 721 (UniProtKB Q5GPYG-3) was mapped to column 745 of the MAFFT alignment by iterating over the gapped human sequence and counting non-gap residues; residue states at this column were extracted for all taxa and used for ancestral-state reconstruction in PastML [77] using MAP prediction under the F81 model.

### Bioinformatic identification of NEK1-ALS missense variants

Prioritisation of genetic variants was performed using summary statistics from a recent exome-wide rare variant association study involving 13,138 ALS patients and 69,775 controls [11]. Variants were annotated with a pass_callrate flag indicating whether they met the stringent call-rate quality control thresholds required for unbiased exome-wide discovery analyses. An additional set of variants failing these thresholds, but present in genes curated by the ALS Gene Curation Expert Panel (GCEP) was also evaluated in the original study, consistent with the rationale that relaxing call-rate filters for established ALS-associated loci enables more comprehensive assessment of known disease genes ([11]; Figure 1b therein). For the purposes of variant prioritisation in the present study, only variants passing the stringent call-rate filters (pass_callrate = TRUE) were considered, and missense variants in NEK1 nominally associated with ALS risk (P < 0.05) were selected for functional characterisation. This threshold was used to prioritise variants with genetic evidence of enrichment for functional follow-up, rather than to define pathogenicity. The complete single-variant association statistics for NEK1, including both pass_callrate = TRUE and FALSE variants, are provided in **electronic supplementary material, Table S6**. Statistical associations with ALS risk were tested at the variant, gene and domain level using Firth logistic regression and the Rare Variant Association Toolkit (RVAT; [78]). All association testing analyses included correction for sex, genome-wide burden of synonymous variants, and the first 10 principal components derived from principal component analysis of common variant profiles. Variant annotations were generated using SnpEff for nonsense and frameshift mutations [75] and splice variants via dbscSNV [80], with annotations based on Ensembl release 105 for build GRCh38 of the human genome.

### Data accessibility

All data needed to evaluate the conclusions in the paper are present in the paper and/or the electronic supplementary material. Primary/source data associated with the main and supplementary figures, sequencing data for CRISPR/Cas9-generated U-2 OS *NEK1* knockout cells, and sequencing data for all plasmids generated or used in this study are publicly deposited in the Zenodo Data Repository (10.5281/zenodo.2211813G).

Annotated peptide spectra shown in the supplementary data are Mascot-derived peptide-spectrum match reports rather than raw mass spectra. The mass spectrometry proteomics data have been publicly deposited to the ProteomeXchange Consortium via the PRIDE [81] partner repository with the dataset identifier PXD075815.

A step-by-step phosphomapping protocol has been submitted, upon invitation, to Bio-protocol. One of the major aims of this work was to create a suite of robust resources for the community; accordingly, all cell lines, reagents and antibodies generated in-house will be made available upon reasonable request from the Corresponding Author, A.R.M.

### Ethics

Custom sheep polyclonal antibodies were generated under UK Home Office Project Licence PP545072G. All animal procedures were carried out in accordance with the Animals (Scientific Procedures) Act 1986 (ASPA), and were ethically reviewed and approved by the University of Edinburgh’s Roslin Animal Welfare and Ethical Review Body (AWERB) via the Named Veterinary Surgeon (NVS), prior to commencement. Animal welfare and the principles of the 3Rs (Replacement, Reduction and Refinement) were considered throughout, and *in vivo* procedures were in accordance with the ARRIVE guidelines.

No new human participants were recruited and no human-derived samples were collected for this study. Human cell lines used were established lines, and human genetic association statistics were obtained from a previously published study [11], conducted with the ethical approvals described therein.

### CRediT (Contributor Roles Taxonomy)

Conceptualisation (S.A., J.R., A.R.M.); Data curation (S.A., R.G., I.S., A.R.M.); Formal analysis (S.A., S.A.A.R., R.G., I.S., A.R.M.); Funding acquisition (A.R.M.); Investigation (S.A., S.A.A.R., I.M.M., A.K., P.J.H., A.R.M.); Methodology (S.A., S.A.A.R., P.J.H., J.H.V., K.P.K., A.R.M.); Project administration (A.R.M.); Resources (F.B., T.M.); Software (S.A.A.R., I.S., A.R.M.); Supervision (A.R.M.); Validation (S.A., A.R.M.); Visualisation (S.A., S.A.A.R., A.R.M.); Writing – original draft (A.R.M.); Writing – review & editing (all authors).

### Funding

A.R.M. is grateful to the WoodNext Foundation, a component fund administered by Greater Houston Community Foundation, for its generous support of his laboratory’s NEK1-ALS research. He also acknowledges MND Translational Accelerator Initiative (MNDAcc) funding from the UK Medical Research Council and National Institute for Health and Care Research, managed by Dementias Platform UK (MR/Z505535/1), and a Royal Society research grant (RGS/R2/252027).

## Acknowledgements

We dedicate this work to the memory of Dr Andrew Johnston Davies, whose experience with NEK1-associated ALS continues to motivate research into the molecular basis of motor neuron disease and its diverse genetic causes. We are grateful to MRC PPU Lab Management (led by: Dr Edwin Allen, Jessica Blackburn and Emma Webster), MRC Reagents and Services (led by Dr James Hastie; Dr Nicola T. Wood’s team for cloning and Hannah Leech for help with virus production), MRC PPU Mass Spectrometry facility (led by Dr Renata Soares), and MRC PPU Operations and Administrative support (led by Dr Monika Zwirek and Alison Hart) for their efficient assistance throughout this project. We acknowledge Dr Manas Sahu for generating the NEK1-ALS mutation plasmids via site-directed mutagenesis using NEB Q5^®^ Site-Directed Mutagenesis Kit when he was in Mehta Lab, and Dr Bethany Geary for assisting with the implementation of Skyline software-based XIC analysis. A.R.M. and S.A.A.R. thank Professor Ronald Hay FRS FRSE FMedSci and Dr Anna Plechanovova in his lab for providing access to their mass photometry instrumentation. A.R.M. thanks Jessica Rietbrock (Proteintech), Robyn Parish (Fisher/Thermo Fisher Scientific), Ellie Fraser (Fisher Scientific), and Dr Leon Williamson (Merck Life Science) for their support with the reagents used in this study. Finally, A.R.M. thanks Professor Sir Philip Cohen FRS FRSE FMedSci, Professor Dario Alessi OBE FRS FRSE FMedSci and Professor Miratul Muqit FMedSci FRSE for helpful scientific discussions.

## Use of Artificial Intelligence (AI) and AI-assisted technologies

Claude Opus 4.8 was used to assist with generation of plotting code and compilation of phosphosite worksheets by combining mass spectrometry output tables into curated summary spreadsheets. All outputs were manually reviewed and verified by the corresponding author against the source data.

## Electronic supplementary material figure legends

**Figure S1:**
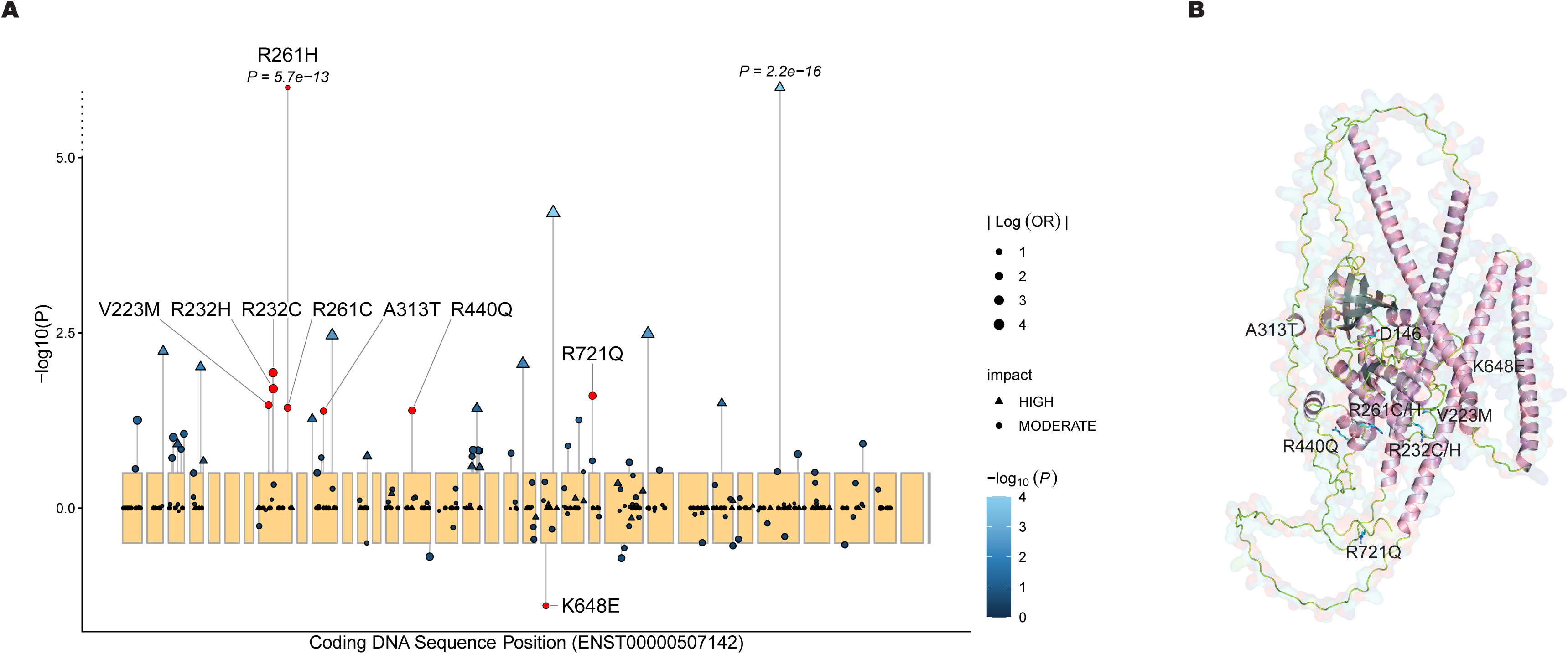
Extended rare variant association statistics and structural mapping of ALS-associated NEK1 missense variants. **(A)** Lollipop plot of association statistics for rare NEK1 variants of moderate or high predicted impact, plotted by coding DNA sequence position (ENST00000507142.6). As in Figure 1B, the plot is restricted to variants passing the call-rate quality-control filters (see Methods) and with a minor allele frequency below 0.05. Each point represents one variant; point size reflects |Log(OR)|, shape indicates predicted mutational impact (triangle: HIGH; circle: MODERATE), and fill colour indicates –log10(*P*) on a scale truncated at 4, chosen so that variation amongst the majority of variants is resolvable. Variants exceeding the plotted y-axis range are shown above an axis break and annotated with their exact *P* value. Variants drawn in red, in place of the – log10(*P*) colour scale, are the nine missense variants selected for functional characterisation in this study, the same nine shown in Figure 1B; note that further variants of high predicted impact also reach P < 0.05 but were not selected. These nine are labelled by amino acid substitution, with leader lines indicating the corresponding data points. Yellow shading demarcates individual exons. Variants orientated below the baseline are less frequent in ALS cases than in matched controls. MODERATE impact variants comprise missense and inframe deletion variants, and HIGH impact variants comprise nonsense and frameshift mutations, all predicted using SnpEff, together with splice acceptor/donor variants predicted using dbscSNV. **(B)** AlphaFold3 structural model of full-length NEK1 with all ALS-associated missense variants selected for functional characterisation indicated at their respective positions (V223M, R232C/H, R261C/H, A313T, R440Q, K648E, R721Q). D146, which when mutated to alanine renders NEK1 kinase-dead, is also indicated as a structural reference point.

**Figure S2:**
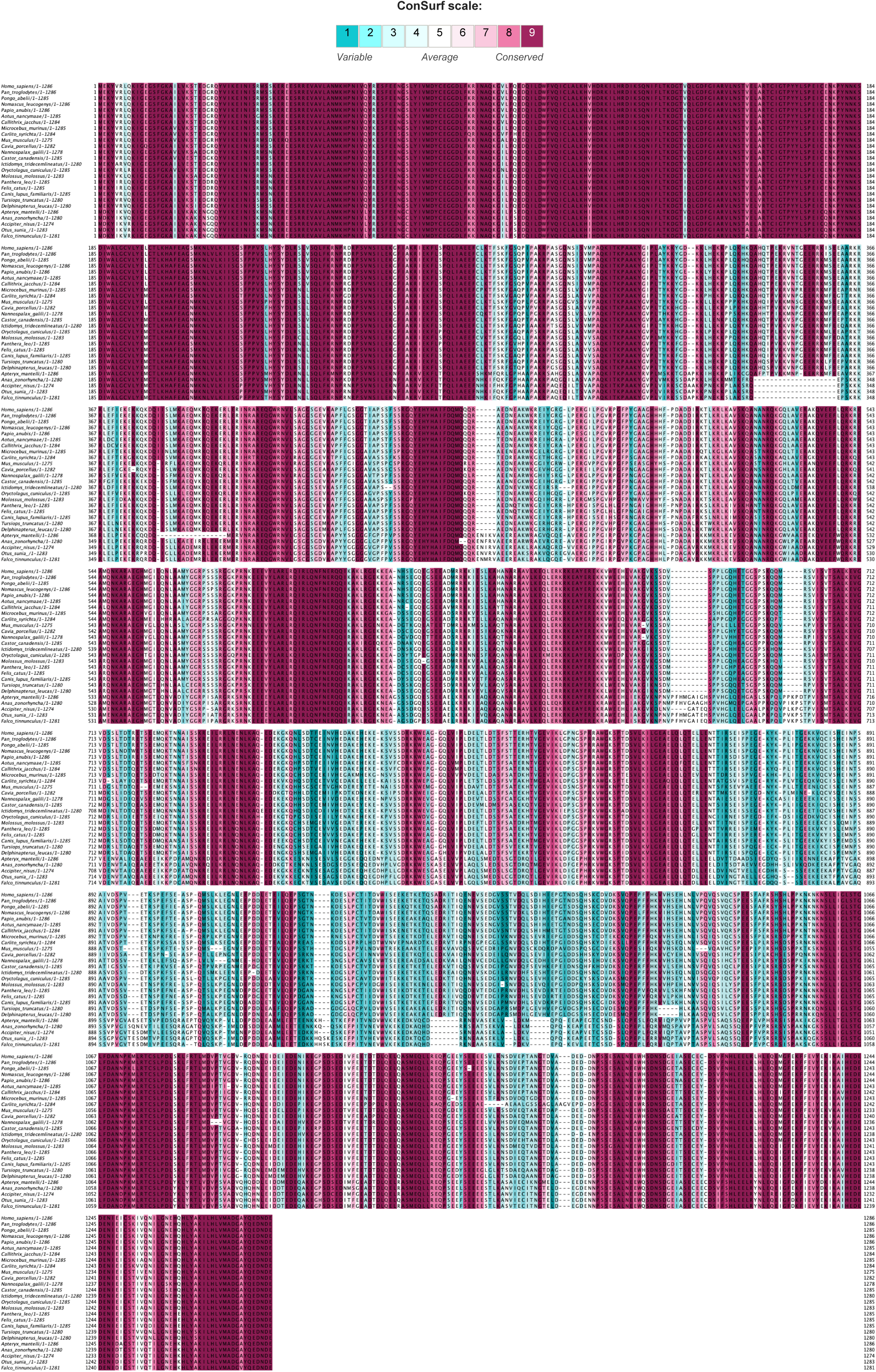
Evolutionary conservation of NEK1 across 26 vertebrate species. Multiple sequence alignment of NEK1 protein sequences from 26 vertebrate species, generated using MAFFT with default parameters. Per-residue evolutionary conservation scores were calculated using ConSurf with human NEK1 isoform 3 (UniProtKB Q5GPYG-3) as the query, and are displayed using the standard nine-colour ConSurf palette (1, most variable; 9, most conserved). Residues in the human sequence corresponding to alignment gaps are uncoloured.

**Figure S3:**
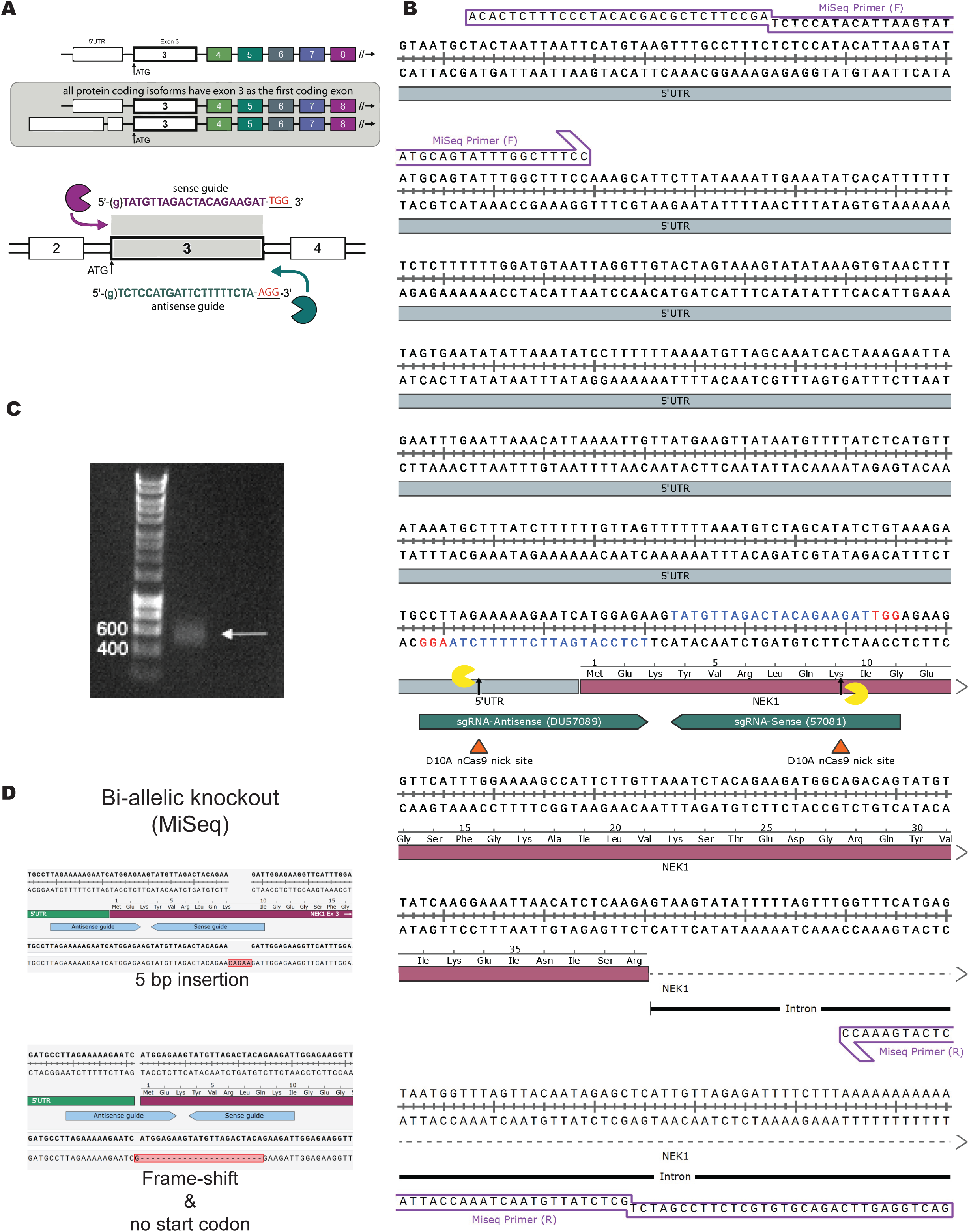
CRISPR-Cas9 nickase-mediated generation and validation of a *NEK1* homozygous knockout U-2 OS cell line. **(A)** Schematic of the CRISPR-Cas9 nickase targeting strategy. Exon 3 was selected as the target because it is the first coding exon shared by all *NEK1* protein-coding isoforms, ensuring complete knockout of all expressed variants. A paired nickase approach using D10A Cas9 was employed, with sense (DU57081) and antisense (DU57089) guide RNAs positioned to nick opposite strands in close proximity to the ATG start codon. **(B)** Annotated sequence map of the *NEK1* exon 3 locus, showing the positions of the sense and antisense sgRNAs, D10A nCas9 nick sites, MiSeq primer binding sites (forward and reverse), the 5′UTR, and the beginning of the *NEK1* coding sequence with translated amino acids indicated. **(C)** 1% Agarose gel showing the 481 bp wild-type PCR amplicon (arrow) spanning the CRISPR target site, used as input for amplicon-based next-generation sequencing. **(D)** MiSeq sequencing confirmation of biallelic *NEK1* gene disruption. Both alleles harbour distinct frameshift-inducing indels: one allele carries a 5 bp insertion introducing a premature stop codon, and the second allele carries a frameshift mutation that abolishes the start codon, confirming complete loss of NEK1 protein expression from both alleles.

**Figure S4:**
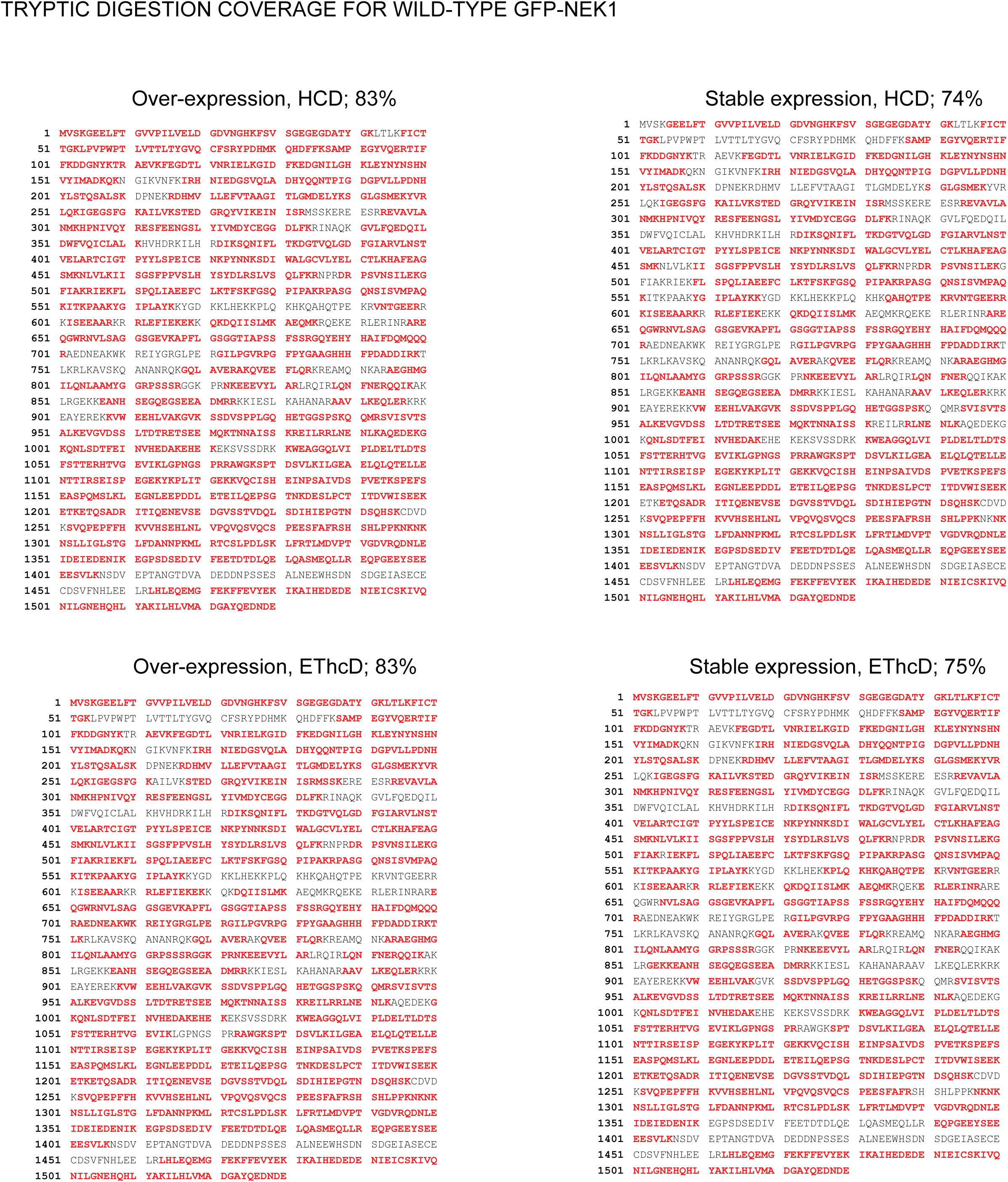
Tryptic digestion sequence coverage of wild-type GFP-NEK1. Sequence coverage maps for wild-type GFP-NEK1 following Trypsin/Lys-C in-gel digestion and LC-MS/MS analysis, shown as a representative example of coverage achieved across all samples. Data are shown for transient overexpression and stable expression conditions using HCD and EThcD fragmentation (83%, 83%, 74%, and 75% coverage respectively). Detected peptides are highlighted in red; undetected regions are shown in black. In these maps EGFP occupies construct residues 1–235, the Ser-Gly-Leu-Gly-Ser linker residues 240–244 and NEK1 isoform 3 residues 245–1530; NEK1 isoform 3 numbering is obtained by subtracting 244.

**Figure S5:**
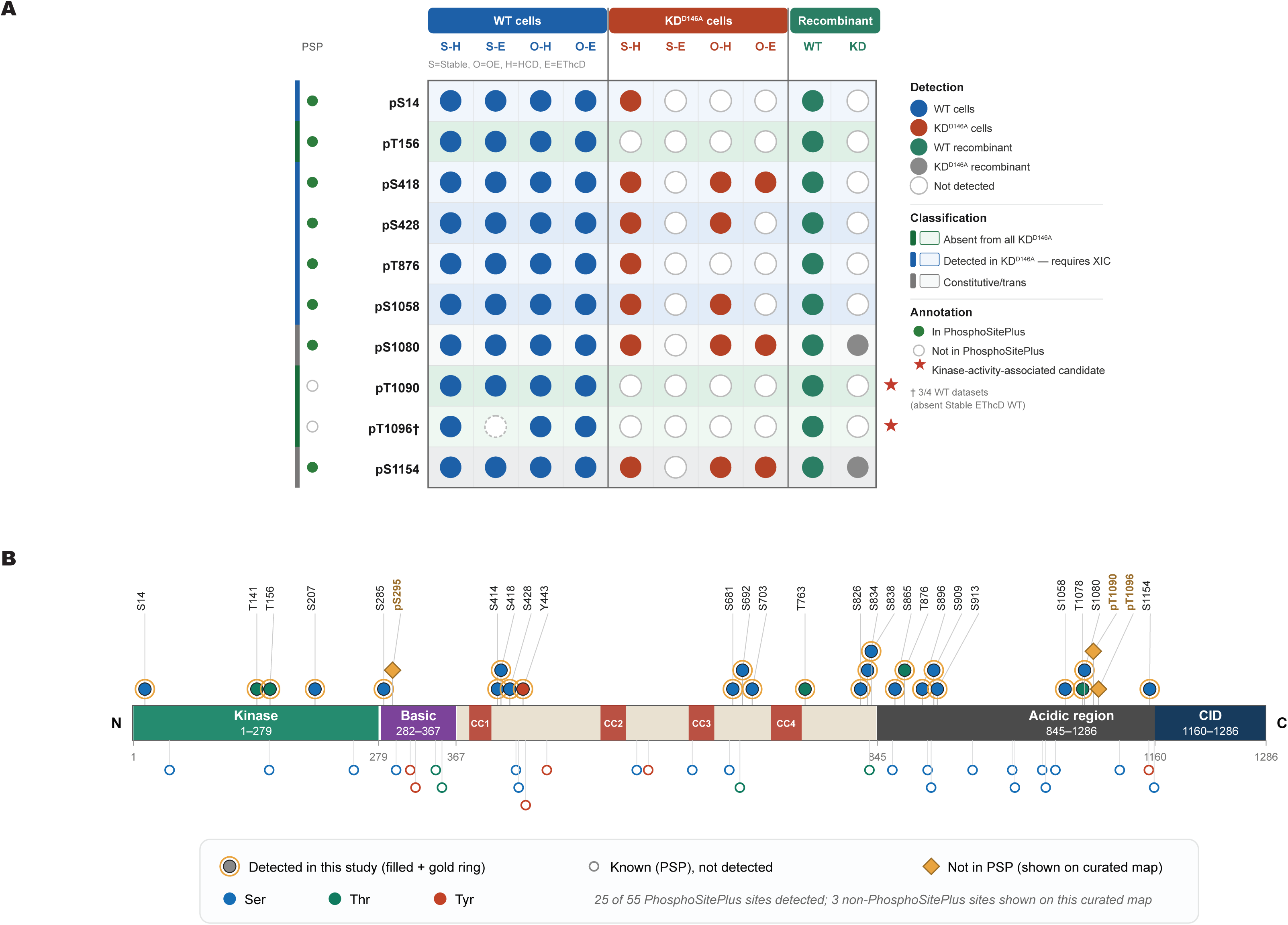
Detection of NEK1 phosphorylation sites across all experimental datasets. (**A**) Matrix summarising detection across the cell-based acquisitions—WT and KD^D14GA^ NEK1, each in the stable and overexpression systems by HCD and EThcD—together with the recombinant WT and KD^D14GA^ datasets. Rows (isoform 3 coordinates) comprise the eight kinase-activity-associated candidate phosphosites carried forward for XIC quantification (pSer14, pThr156, pSer418, pSer428, pThr876, pSer1058, pThr1050 and pThr105G), together with the two recurrent sites detected in recombinant KD^D14GA^ (pSer1080 and pSer1154; classified as constitutive/trans-phosphorylated). Filled symbols denote detection and open symbols non-detection in each dataset; the left-hand column indicates annotation in PhosphoSitePlus^®^ (filled, annotated; open, not annotated). Colour banding classifies each site as absent from kinase-dead datasets, detected in KD^D14GA^ datasets and therefore requiring XIC quantification, or constitutive/trans-phosphorylated. Sites prioritised as kinase-activity-associated candidates (pThr1050 and pThr105G) are starred; pThr1096 was detected in three of the four WT cellular datasets (absent from the Stable EThcD dataset). (**B**) All 55 PhosphoSitePlus^®^-annotated phosphorylation sites mapped onto the NEK1 isoform 3 domain architecture (UniProtKB Q5GPYG-3; 1,286 aa). Sites detected in this study are shown above the protein bar as filled circles with a gold ring (*n* = 25); known PhosphoSitePlus^®^ sites not detected are shown below as open circles (*n* = 30). Three selected non-PhosphoSitePlus^®^ sites are indicated by gold diamonds on this curated map: pSer255, pThr1050 and pThr105G. Additional non-PhosphoSitePlus^®^ detections present in subset datasets are listed in Table S1. Residue type is colour-coded (blue, Ser; green, Thr; red, Tyr). Dark red segments within the linker region denote the four predicted coiled-coil domains (CC1, 382–407; CC2, 531–560; CC3, G31–GG0; CC4, 724–759).

**Figure S6:**
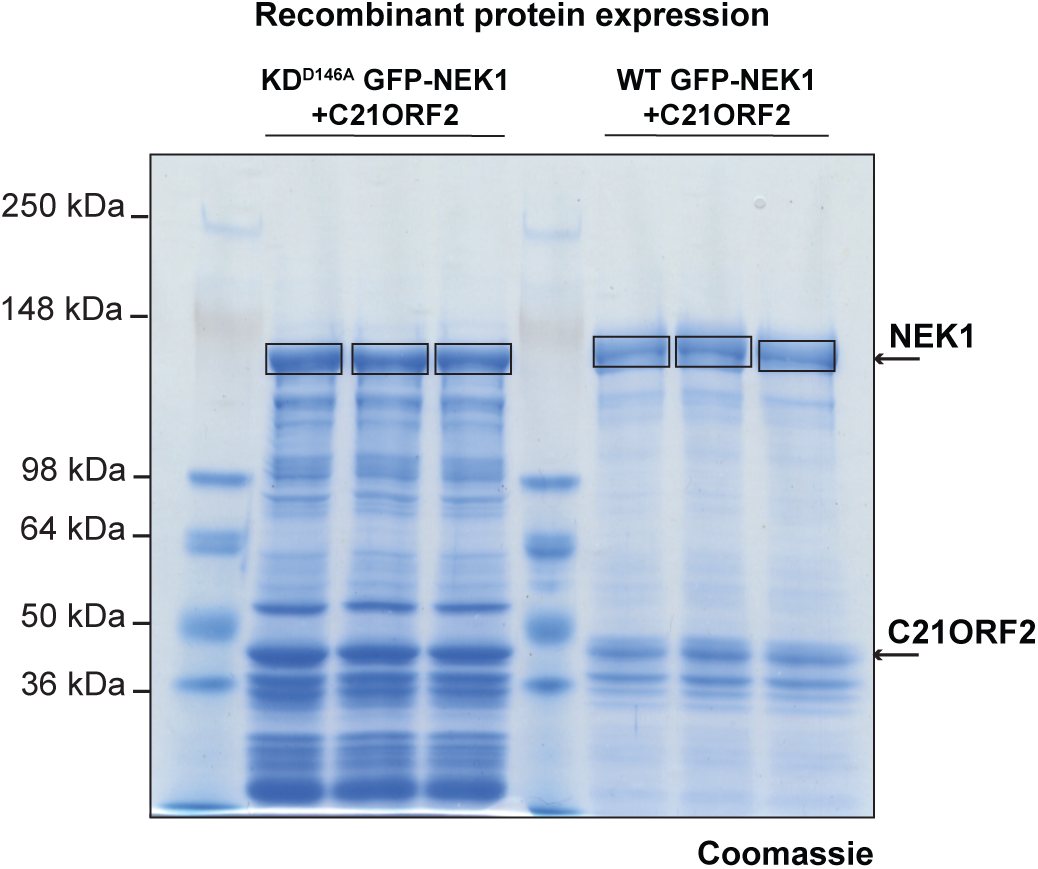
Recombinant expression and purification of NEK1 and the NEK1–C21ORF2 complex. Coomassie Brilliant Blue-stained SDS-PAGE of WT and KD^D14GA^ NEK1, co-expressed with C21ORF2 in Sf9 insect cells as GFP-tagged constructs, purified by anti-GFP affinity chromatography and recovered by RV-3C cleavage before SDS-PAGE (three independent preparations per construct). Bands corresponding to NEK1 (boxed) and C21ORF2 are indicated; the NEK1 bands were excised for in-gel digestion and LC-MS/MS analysis. Molecular-weight markers (kDa) are shown at left.

**Figure S7:**
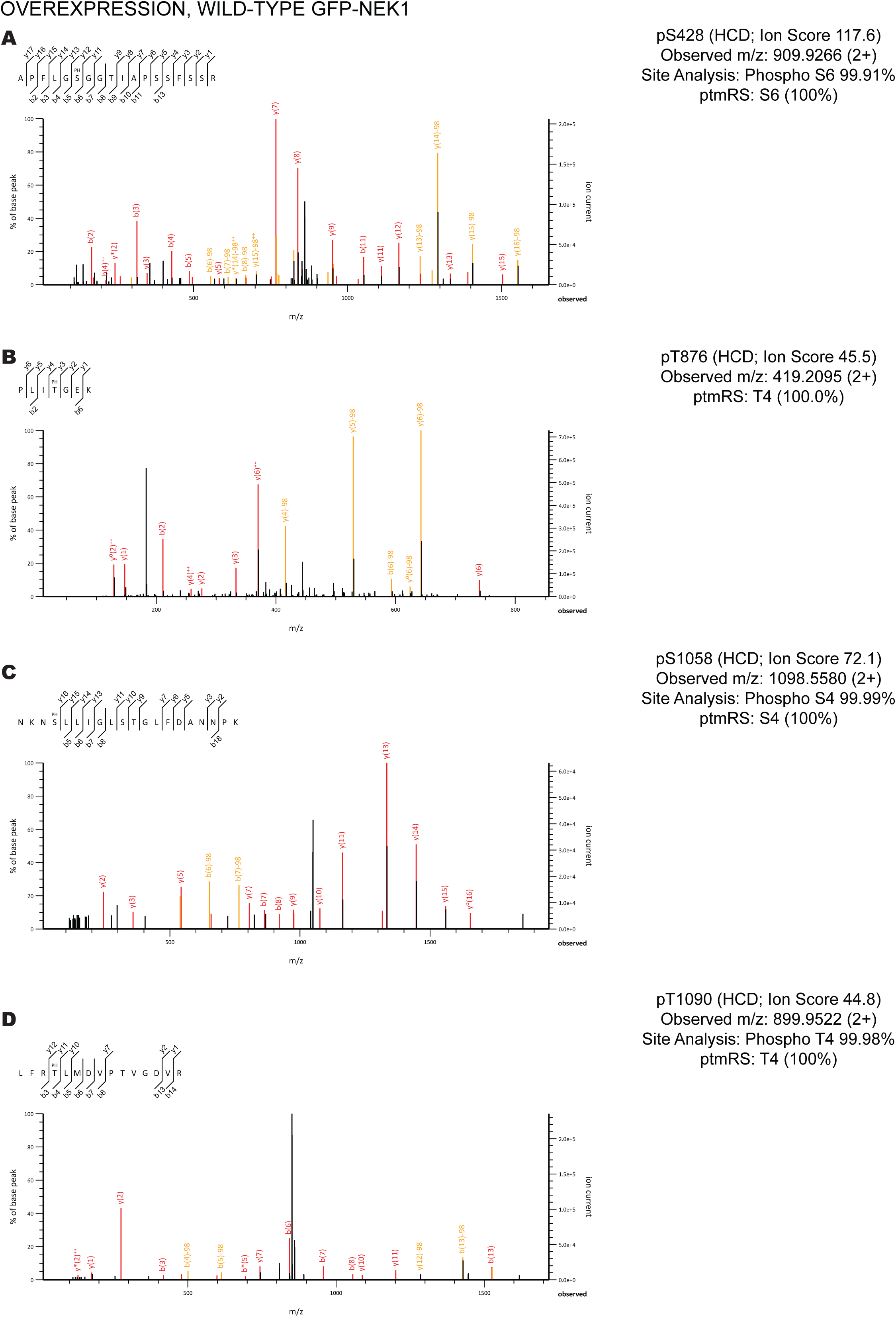
Mascot-derived annotated peptide-spectrum match reports supporting phosphosite localisation for pSer428, pThr876, pSer1058 and pThr1090 in wild-type GFP-NEK1 following transient overexpression. Representative Mascot-derived annotated HCD MS/MS spectra from transient overexpression of wild-type GFP-NEK1 in *NEK1* knockout U-2 OS cells. Sequence ladders above each spectrum indicate matched backbone cleavages, with the phosphorylated residue marked PH. Sequence ions (b- and y-series) are shown in red; phosphorylation-directed neutral loss ions (−H₃PO₄, −98 Da) are shown in orange; unassigned peaks are shown in black. Positional annotations refer to residue numbering within the tryptic peptide; NEK1 isoform 3 site identities are given in parentheses. **(A)** pSer428, identified by HCD fragmentation of tryptic peptide APFLGSGGTIAPSSFSSR (Ion Score 117.6; observed m/z 505.52GG, [M+2H]²⁺; Mascot site analysis: Phospho SG 55.51%; ptmRS: S6 100%). **(B)** pThr876, identified by HCD fragmentation of tryptic peptide PLITGEK (Ion Score 45.5; observed m/z 415.2055, [M+2H]²⁺; ptmRS: T4 100.0%). Thr4 is the sole phosphorylatable residue within this peptide; site localisation is therefore unambiguous by definition. The prominent unlabelled peak at m/z 183.15 corresponds to the a(2) ion (PL; b2 − CO), a characteristic N-terminal fragment not annotated by default in Mascot. **(C)** pSer1058, identified by HCD fragmentation of tryptic peptide NKNSLLIGLSTGLFDANNPK (Ion Score 72.1; observed m/z 1098.5580, [M+2H]²⁺; Mascot site analysis: Phospho S4 55.55%; ptmRS: S4 100%). **(D)** pThr1090, identified by HCD fragmentation of tryptic peptide LFRTLMDVPTVGDVR (Ion Score 44.8; observed m/z 855.5522, [M+2H]²⁺; Mascot site analysis: Phospho T4 55.58%; ptmRS: T4 100%). The dominant unlabelled peak at m/z 855.55 corresponds to the unfragmented doubly-charged precursor ion.

**Figure S8:**
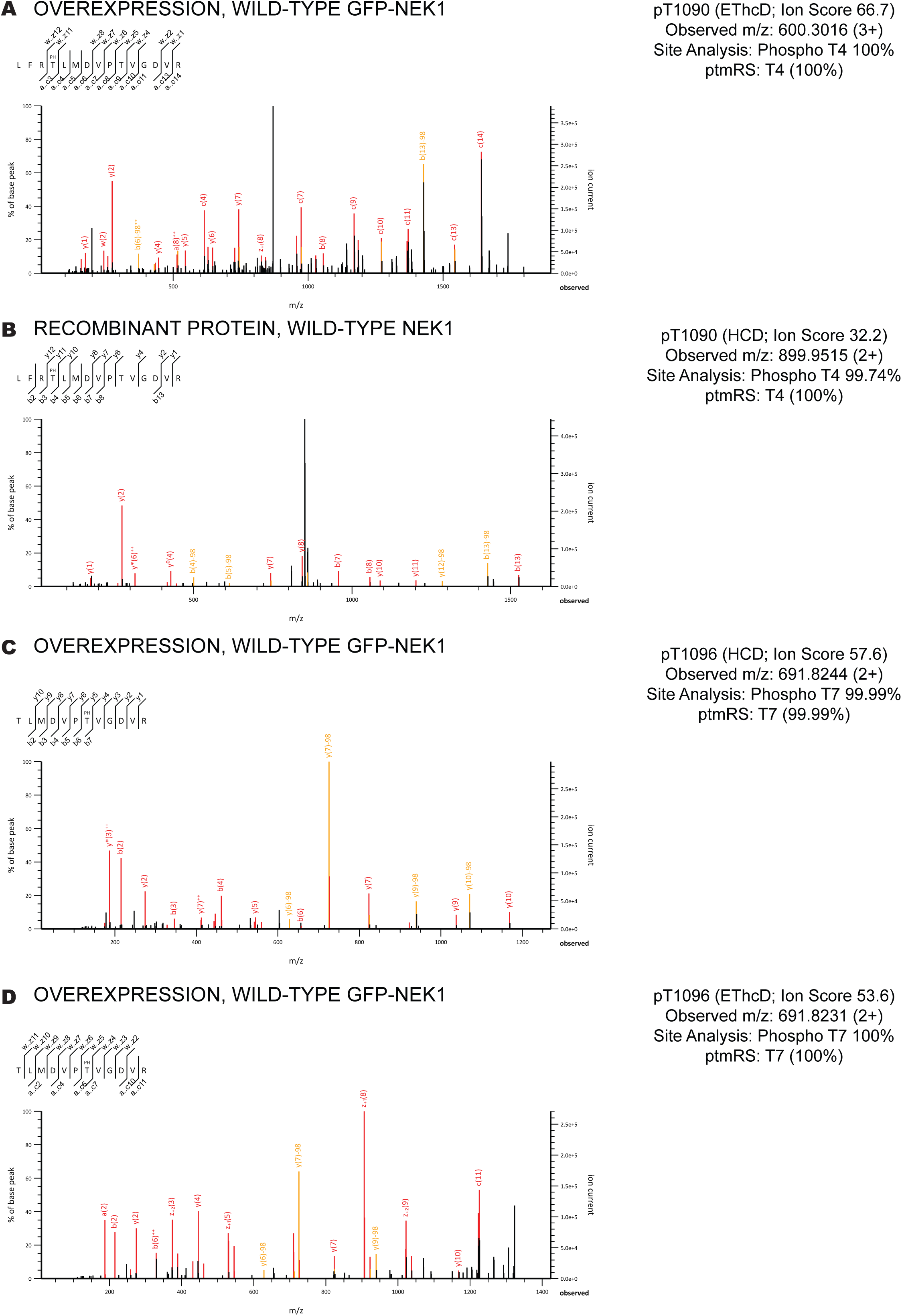
Mascot-derived annotated peptide-spectrum match reports supporting phosphosite localisation for pThr1090, detected by EThcD and recombinant HCD, and pThr1096, detected by HCD and EThcD, in wild-type cell-based GFP-NEK1 and recombinant NEK1 datasets. Annotated Mascot-derived MS/MS spectra confirming phosphosite localisation for pThr1050 and pThr105G. **(A, D)** EThcD panels: c-ions and z- radical ions arising from the electron-transfer component are shown in red alongside HCD-derived b/y-ions; sequence ladder tick marks denote matched a/c-ions (below sequence) and w/z-ions (above sequence) as annotated by Mascot for EThcD data. **(B, C)** HCD panels: sequence ions (b- and y-series) are shown in red; phosphorylation-directed neutral loss ions (−H₃PO₄, −98 Da) are shown in orange. Unassigned peaks are shown in black throughout. Positional annotations refer to residue numbering within the tryptic peptide. **(A)** pThr1090, confirmed by EThcD fragmentation of tryptic peptide LFRTLMDVPTVGDVR from transient overexpression of WT GFP-NEK1 in *NEK1* knockout U-2 OS cells (Ion Score 66.7; observed m/z G00.301G, [M+3H]³⁺; Mascot site analysis: Phospho T4 100%; ptmRS: T4 100%). The dominant unlabelled peak at m/z ∼900 corresponds to the charge-reduced precursor ([M+2H]²⁺•), a characteristic artefact of electron-transfer fragmentation. **(B)** pThr1090, confirmed by HCD fragmentation of tryptic peptide LFRTLMDVPTVGDVR from recombinant WT NEK1 purified from Sf9 insect cells after GFP-affinity purification and RV-3C tag cleavage (Ion Score 32.2; observed m/z 855.5515, [M+2H]²⁺; Mascot site analysis: Phospho T4 99.74%; ptmRS: T4 100%). This spectrum provides independent, cell-free support for pThr1090 detection in WT recombinant NEK1. **(C)** pThr1096, identified by HCD fragmentation of tryptic peptide TLMDVPTVGDVR from transient overexpression of WT GFP-NEK1 in *NEK1* knockout U-2 OS cells (Ion Score 57.6; observed m/z 691.8244, [M+2H]²⁺; Mascot site analysis: Phospho T7 55.55%; ptmRS: T7 55.55%). Note that this peptide spans both Thr1090 (T1) and Thr1096 (T7); the spectrum unambiguously localises phosphorylation to Thr1096 (T7, 55.55%), with Thr1090 unphosphorylated in this PSM. **(D)** pThr1096, confirmed by EThcD fragmentation of tryptic peptide TLMDVPTVGDVR from transient overexpression of WT GFP-NEK1 in *NEK1* knockout U-2 OS cells (Ion Score 53.6; observed m/z 691.8231, [M+2H]²⁺; Mascot site analysis: Phospho T7 100%; ptmRS: T7 100%).

**Figure S9:**
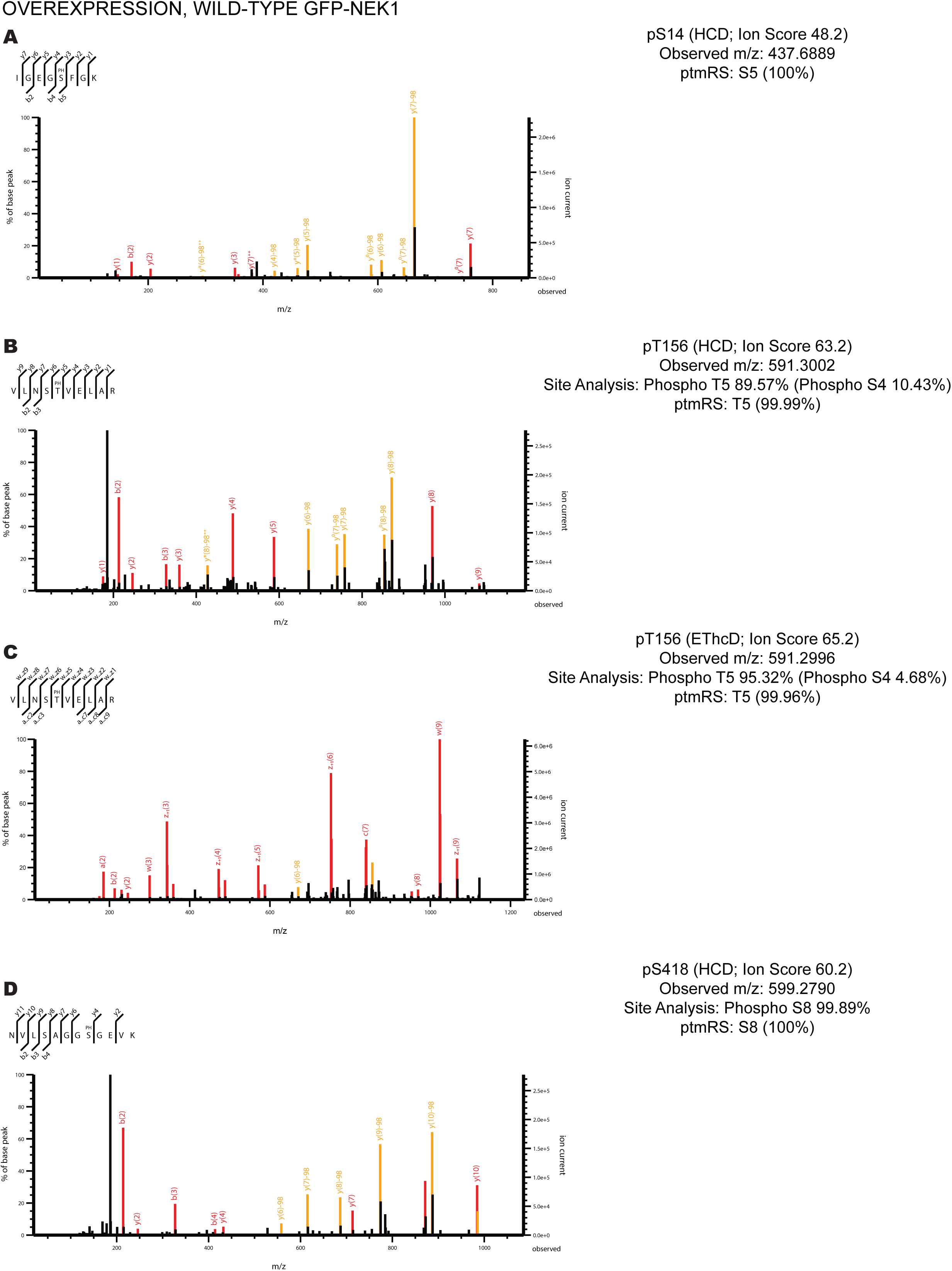
Mascot-derived annotated peptide-spectrum match reports supporting phosphosite localisation for pSer14, pThr156 and pSer418 in wild-type GFP-NEK1—transient overexpression. Representative Mascot-derived annotated MS/MS spectra from transient overexpression of WT GFP-NEK1. **(A)** pSer14, identified by HCD fragmentation (Ion Score 48.2, observed m/z 437.6889, ptmRS: S5 100%). (**B**) pThr156, identified by HCD fragmentation **(Ion Score 63.2**, observed m/z 591.3002, Phospho T5 89.57%, ptmRS: T5 55.55%). (**C**) pThr156, identified by EThcD fragmentation (Ion Score 65.2, observed m/z **551.2556**, Phospho T5 95.32%, ptmRS: T5 55.5G%). **(D)** pSer418, identified by HCD fragmentation (Ion Score 60.2, observed m/z 555.2750, Phospho S8 55.85%, ptmRS: S8 100%). Fragment ions are annotated with b-ions (red) and y-ions (orange); the observed spectrum is shown in black alongside the theoretical spectrum.

**Figure S10:**
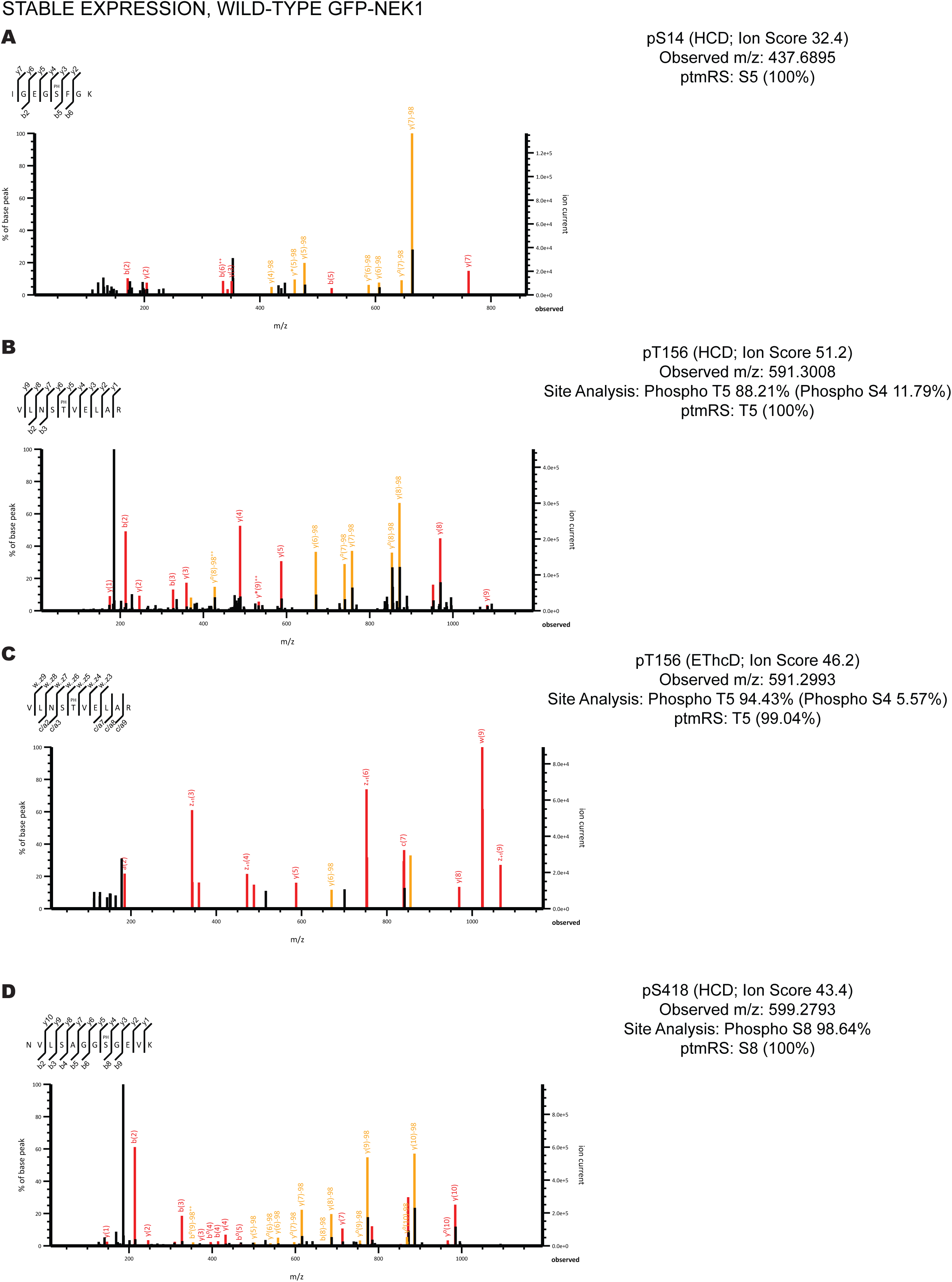
Mascot-derived annotated peptide-spectrum match reports supporting phosphosite localisation for pSer14, pThr156 and pSer418 in wild-type GFP-NEK1—stable expression. Representative Mascot-derived annotated MS/MS spectra from stable doxycycline-inducible expression of WT GFP-NEK1. **(A)** pSer14, identified by HCD fragmentation (Ion Score 32.4, observed m/z 437.6895, ptmRS: S5 100%). **(B)** pThr156, identified by HCD fragmentation (Ion Score 51.2, observed m/z 591.3008, Phospho T5 88.21%, ptmRS: T5 100%). **(C)** pThr156, identified by EThcD fragmentation (Ion Score 46.2, observed m/z 551.2553, Phospho T5 94.43%, ptmRS: T5 99.04%). **(D)** pSer418, identified by HCD fragmentation (Ion Score 43.4, observed m/z 555.2753, Phospho S8 58.G4%, ptmRS: S8 100%). Fragment ions are annotated with b-ions (red) and y-ions (orange); the observed spectrum is shown in black alongside the theoretical spectrum.

**Figure S11:**
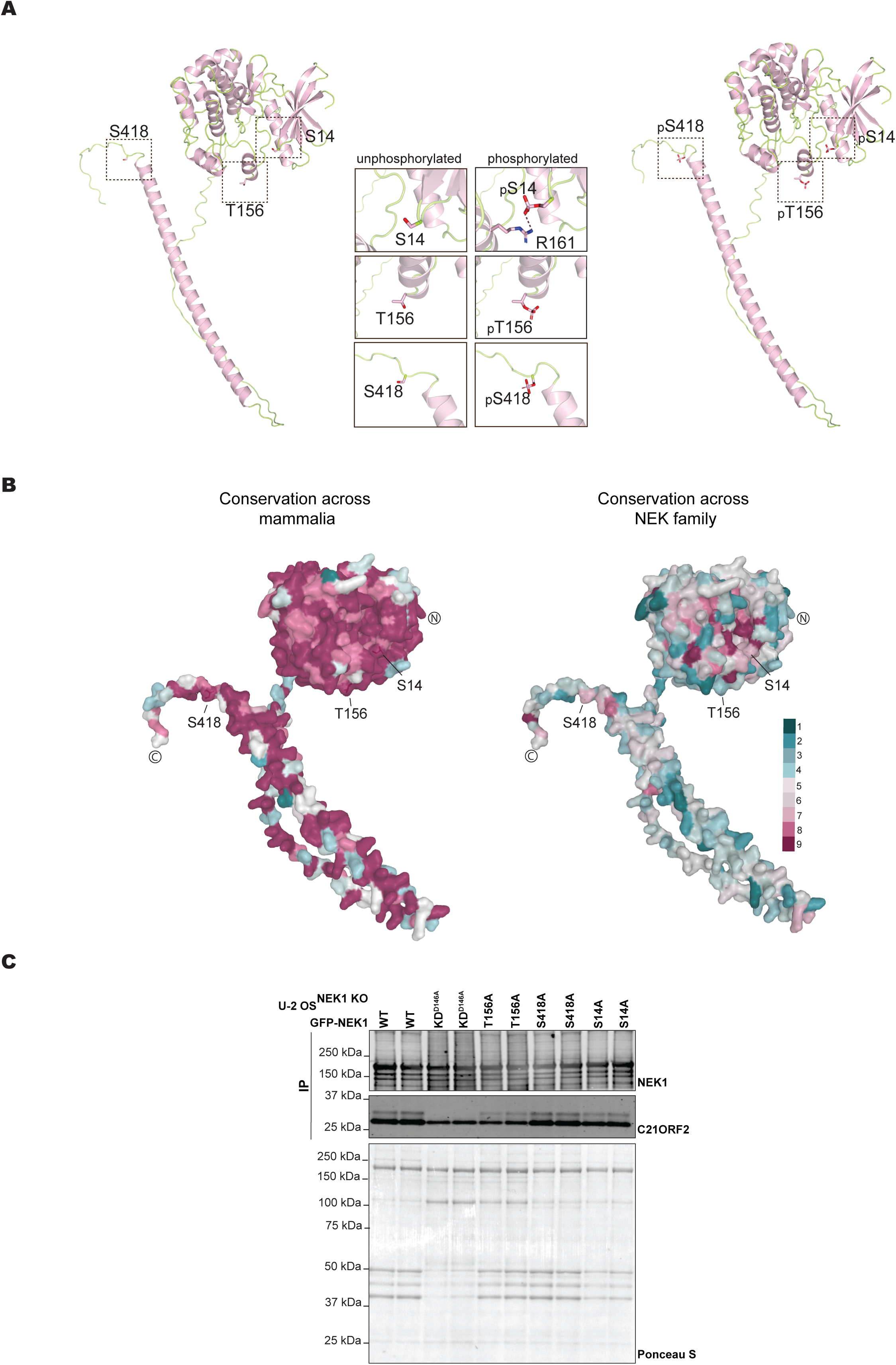
Structural context and conservation of NEK1 autophosphorylation sites, and C21ORF2 binding by phosphosite mutants. **(A)** Predicted structural context of the three NEK1 autophosphorylation sites before and after phosphorylation. AlphaFold3 models of WT NEK1 (residues 1–1,286), with residues 1–431 displayed, showing Ser14, Thr156 and Ser418 in the unphosphorylated state (left) and following phosphorylation (right); in each, the kinase domain is shown at the top and the extended C-terminal coiled-coil below. Boxed regions are enlarged in the central column, unphosphorylated (left) and phosphorylated (right) for each site. Ser14 lies within the glycine-rich (G-) loop of the kinase N-lobe, a region that is disordered and therefore unresolved in the available crystal structure, so its side-chain environment can only be assessed from the model. In the unphosphorylated state, the Ser14 side chain is oriented inwards, towards the body of the N-lobe, whereas Thr156 and Ser418 are solvent-exposed; none of the three residues makes a notable intramolecular contact. The inward orientation of Ser14 should, however, be interpreted with caution: the G-loop is highly mobile, so Ser14 may sample solvent-exposed positions or adopt alternative side-chain rotamers not captured by a single static model. Following phosphorylation, pSer14 is positioned to form an ionic interaction with the guanidinium group of Arg161 (dashed line), whereas pThr156 and pSer418 do not appear to engage in any intramolecular interaction and remain largely solvent-exposed. The engagement of pSer14 by Arg161 suggests that this phosphosite may be partially occluded once formed, which could limit the accessibility of the epitope to phosphospecific antibodies raised against pSer14. **(B)** Surface representation of the same region (residues 1–431) coloured by ConSurf conservation grade (1–9; variable to conserved) across mammalian NEK1 orthologues (left) and across the human NEK kinase family (right), with the N-terminus and the end of the displayed region (residue 431) indicated. All three sites are surface-accessible and highly conserved within the NEK1 lineage, but show more variable conservation across the broader NEK family. **(C)** Anti- GFP co-immunoprecipitation from NEK1 knockout U-2 OS cells expressing WT, KD^D14GA^, or phosphosite-mutant (T156A, S418A, S14A) GFP-NEK1, immunoblotted for NEK1 and C21ORF2. C21ORF2 co-precipitates with NEK1 in all constructs, indicating that the phosphosite substitutions do not disrupt the NEK1– C21ORF2 interaction. A NEK1-dependent C21ORF2 mobility shift is present in WT and in the phosphosite-mutant constructs, but is absent in KD^D14GA^. Ponceau S staining confirms comparable protein loading.

**Figure S12:**
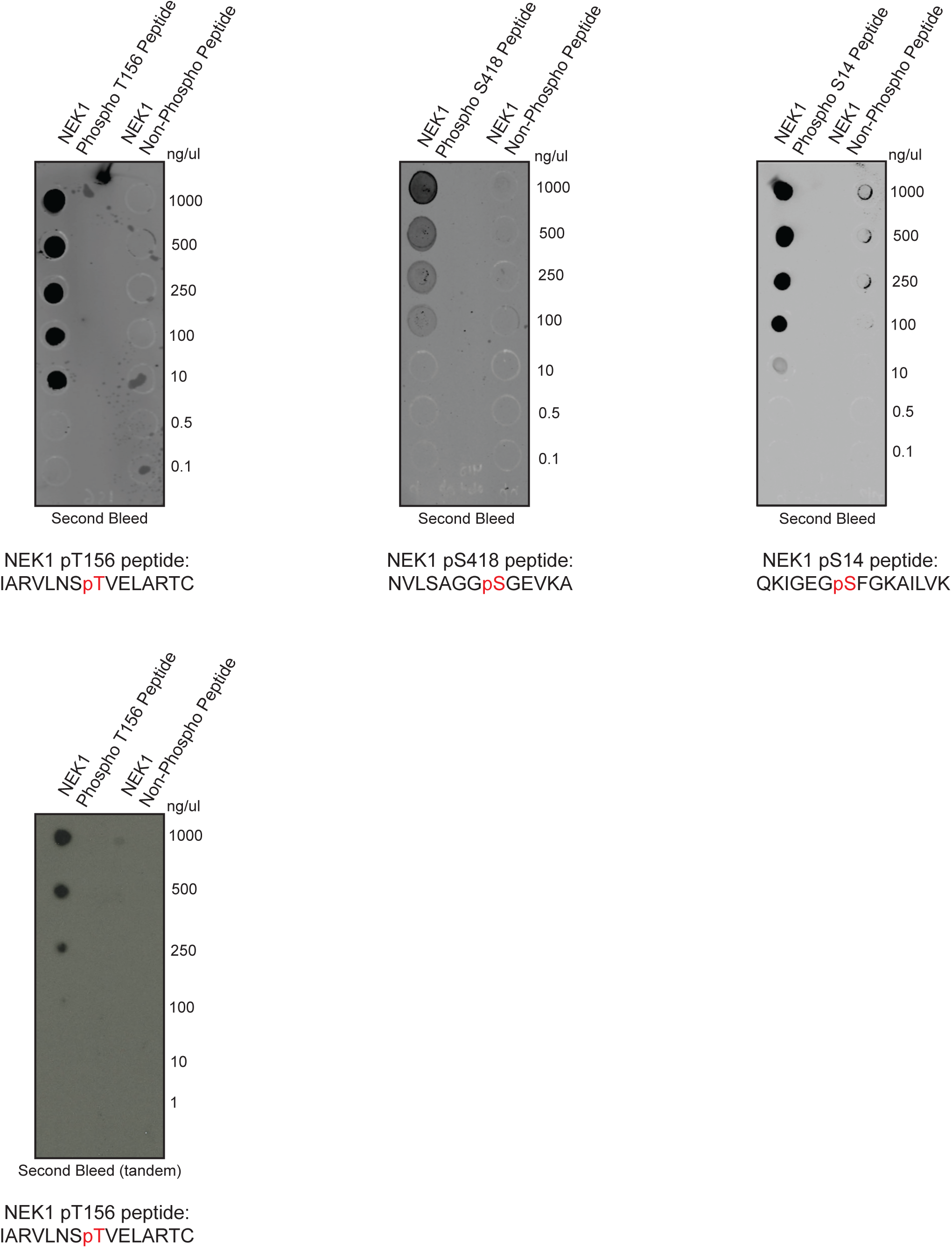
Dot blot validation of phosphospecific antibodies against NEK1 pSer14, pThr156 and pSer418. **(A)** Dot blot analysis of second bleed affinity-purified sheep polyclonal antibodies against pThr156 (upper left), pSer418 (upper middle), and pSer14 (upper right). Serial dilutions of phosphorylated peptide (1000, 500, 250, 100, 10, 0.5, and 0.1 ng/µl), and the corresponding non-phosphorylated peptide counterpart were spotted onto nitrocellulose and probed with each antibody. All three antibodies detect their cognate phosphopeptide in a concentration-dependent manner, with no signal against the non-phospho peptide, confirming phosphospecificity. The pThr156 antibody was additionally subjected to tandem depletion purification (second bleed, tandem; lower panel), in which serum was first passed over non-phosphopeptide resin to remove antibodies recognising the peptide backbone independently of phosphorylation. The tandem-depleted preparation detects the cognate phosphopeptide to 250 ng/µl, compared with 10 ng/µl for the single-step preparation; this reduction in titre is the expected cost of the depletion step and is acceptable at the working concentrations used. The upper and lower panels were acquired as separate experiments, with different exposure settings and slightly different dilution series, and should not be read as a matched side-by-side comparison of sensitivity. The tandem-depleted preparation was used for the immunoblotting experiments reported here. Phosphorylated residues are indicated in red in the peptide sequences shown below each blot.

**Figure S13:**
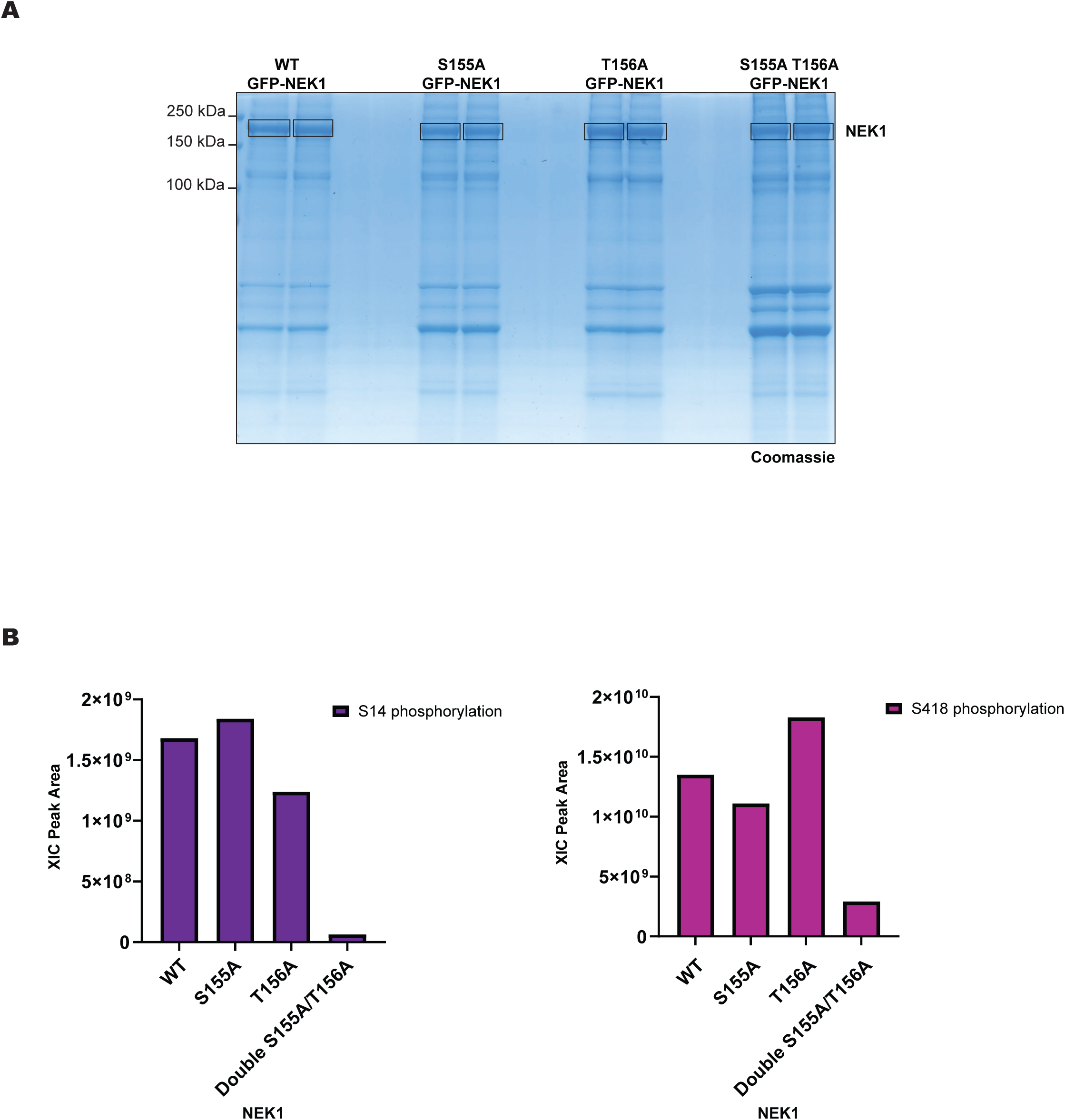
Activation-loop mutagenesis reveals coupling between the Ser155/Thr156 region and distal NEK1 autophosphorylation sites. (**A**) Colloidal Blue-stained SDS-PAGE gel following anti-GFP immunoprecipitation from *NEK1* knockout U-2 OS cells transiently overexpressing WT, S155A, T156A or S155A/T156A GFP-NEK1. Excised NEK1 bands were subjected to in-gel Trypsin/Lys-C digestion and LC-MS/MS analysis. (**B**) XIC peak-area quantification for pSer14 and pSer418 across the indicated GFP-NEK1 constructs, normalised to total ion current within each run. S155A has no appreciable effect on pSer14 or pSer418 relative to WT NEK1, supporting the conclusion that Ser155 is not responsible for the signal assigned to pThr156. The S155A/T156A double mutant causes marked loss of pSer14, an effect not recapitulated by either single mutant, consistent with coupling between pSer14 and the Ser155/Thr156 activation-loop region rather than a strict requirement for Thr156 phosphorylation alone. T156A increases pSer418 relative to WT NEK1, whereas the S155A/T156A double mutant reduces pSer418 relative to T156A, suggesting altered distribution of autophosphorylation within the mutant proteins.

**Figure S14:**
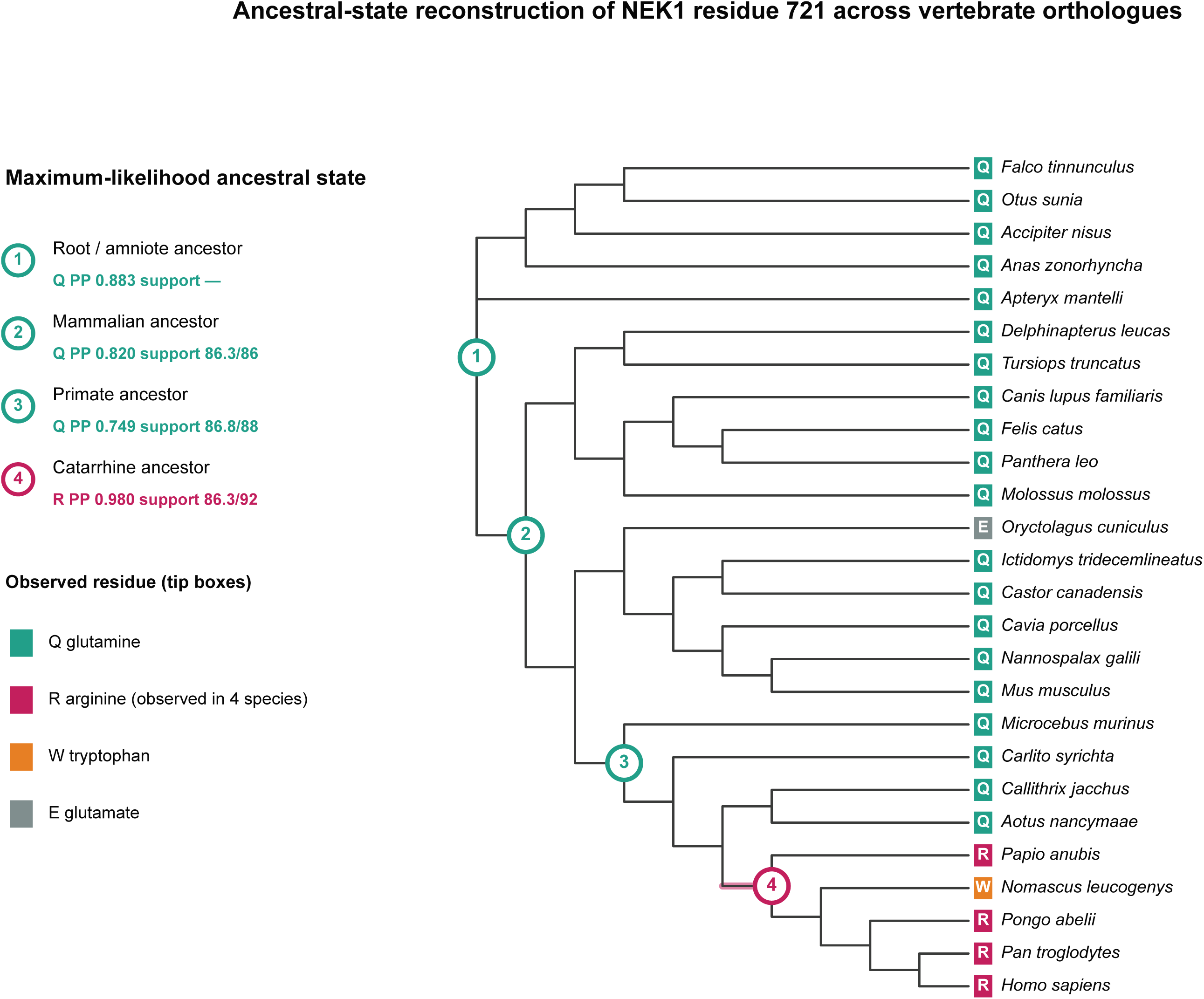
Ancestral-state reconstruction of NEK1 residue 721 across vertebrate orthologues. A maximum-likelihood phylogeny was inferred from a MAFFT alignment of 26 vertebrate NEK1 protein sequences using IQ-TREE v3.0.1 via the CIPRES Science Gateway. ModelFinder selected Q.BIRD+I+G4 as the best-fit model according to BIC, and branch support was assessed using 1000 SH-aLRT and 1000 ultrafast bootstrap replicates. The tree was rooted using avian taxa as the outgroup. Human NEK1 isoform 3 residue 721, corresponding to UniProtKB Q5GPYG-3 and alignment column 745, was analysed using PastML with MAP prediction under the F81 model. Tip boxes indicate observed residues; numbered internal nodes indicate highlighted ancestral-state reconstructions with marginal posterior probabilities, with node support shown as SH-aLRT/UFBoot. The branch on which arginine was inferred to arise is highlighted in magenta. Glutamine (Q) was inferred at the root/amniote, mammalian and primate ancestral nodes, whereas arginine (R) was inferred at the catarrhine ancestor, consistent with a catarrhine-lineage gain of R followed by a gibbon-specific W state. Topology is shown for clarity; branch lengths are not shown to scale. The human R721Q substitution is therefore consistent with reversion toward the inferred ancestral glutamine state.

**Figure S15:**
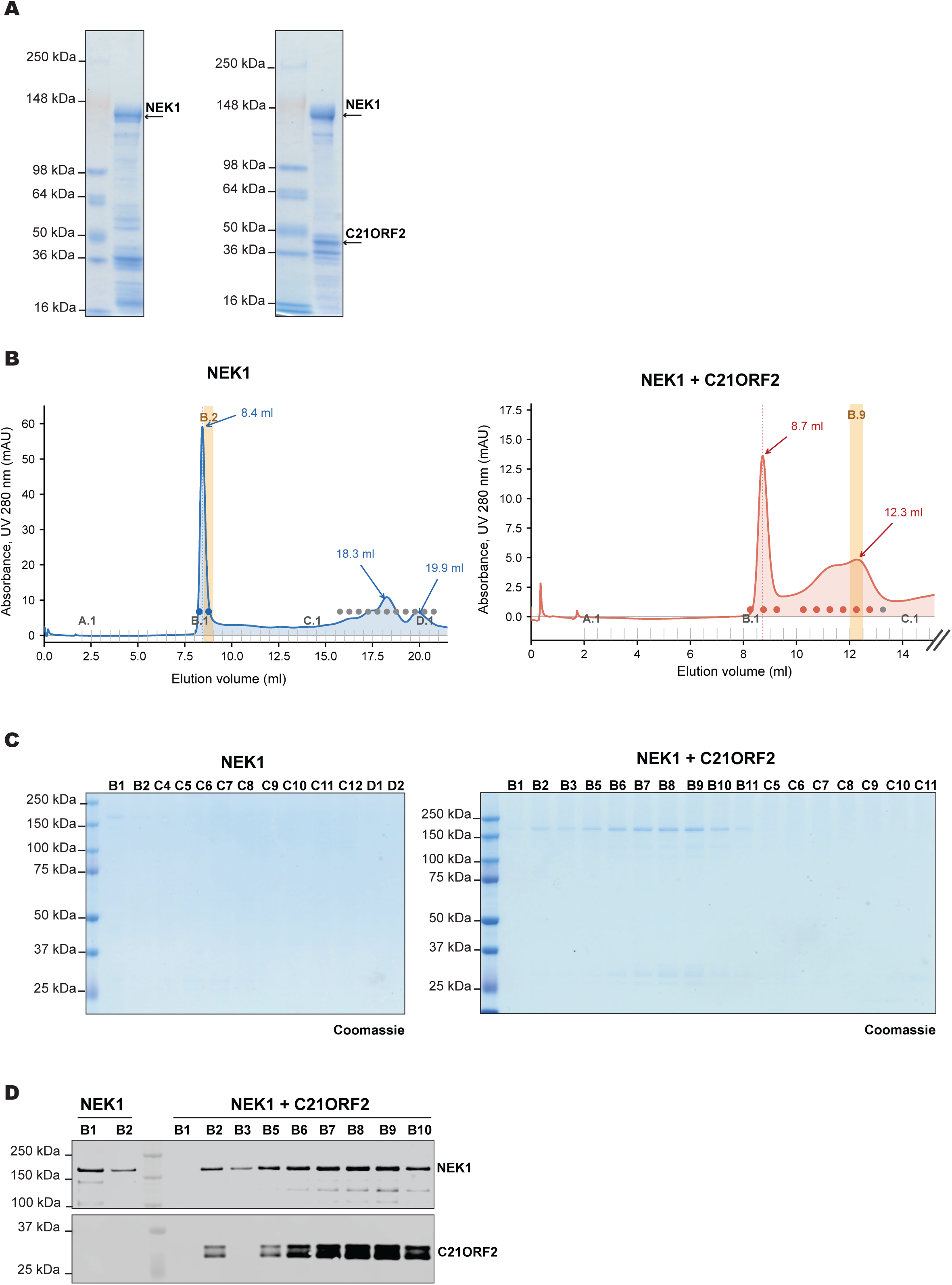
Size-exclusion chromatography of NEK1 and the NEK1– C21ORF2 complex. **(A)** Coomassie-stained SDS-PAGE of purified NEK1 alone and NEK1 co-purified with C21ORF2, with the NEK1 and C21ORF2 bands indicated. **(B)** Size-exclusion chromatograms (Superose 6 Increase 10/300) of NEK1 alone (left) and the NEK1–C21ORF2 complex (right); the void volume for this column is ∼8 ml, and the fractions selected for mass photometry (NEK1 alone, fraction B2, eluting at 8.4 ml; NEK1–C21ORF2, fraction B9, eluting at 12.3 ml) are indicated. **(C)** SDS-PAGE and **(D)** anti-NEK1 and anti-C21ORF2 immunoblot analysis of the collected fractions, confirming co-elution of NEK1 and C21ORF2.

**Figure S16:**
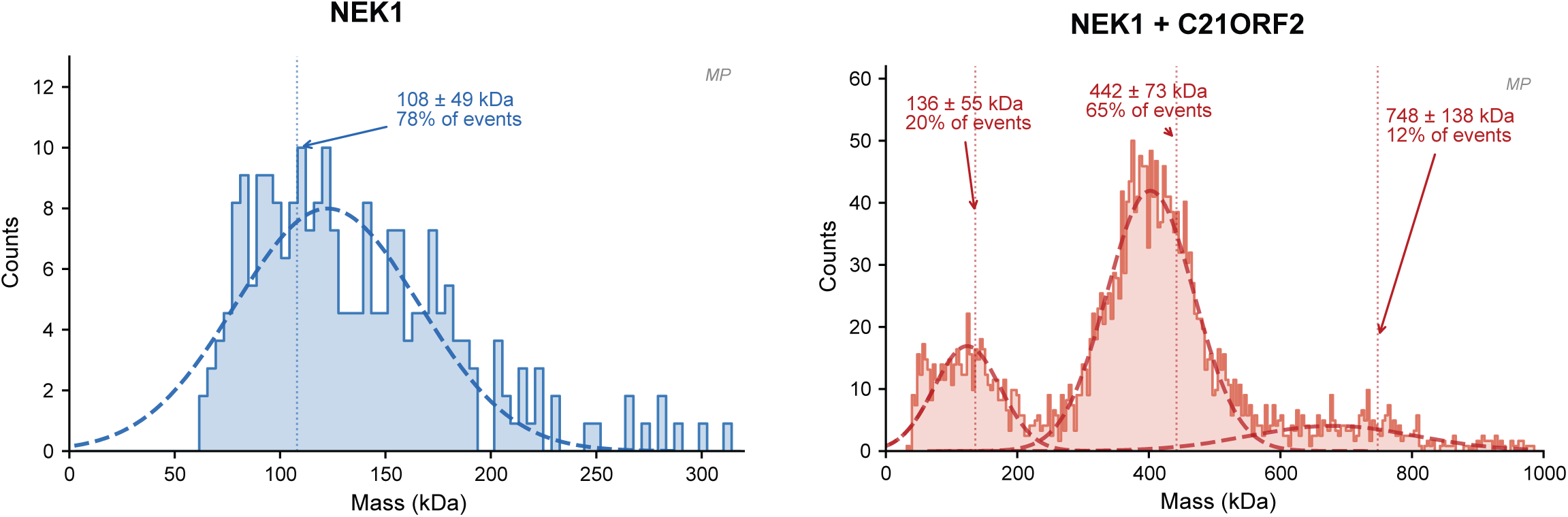
Mass photometry of NEK1 and the NEK1– C21ORF2 complex. Mass distributions measured by mass photometry and fitted with Gaussian functions. NEK1 alone (left) resolves as a predominant species of 108 ± 49 kDa (78% of events), consistent with a monomer; the NEK1– C21ORF2 complex (right) resolves as a major species of 442 ± 73 kDa (65% of events), consistent with a putative 2:4 NEK1– C21ORF2 assembly, together with minor populations of 136 ± 55 kDa (20%) and 748 ± 138 kDa (12%). We note that, in size-exclusion chromatography, NEK1 alone elutes far earlier than expected for a NEK1 monomer (Figure S15B), giving an apparent molecular weight well above the mass measured here. Because mass photometry reports mass independently of shape, this discrepancy is most parsimoniously explained by a non-globular, extended conformation and/or self-association of NEK1 in the absence of C21ORF2. For the complex, the two measurements are concordant: it elutes well within the fractionation range of the column, and its measured mass approximates that calculated for a 2:4 NEK1– C21ORF2 assembly, consistent with a more compact species.

---

**Table S1: Cell-based phosphoproteomic map of wild-type NEK1.** Phosphorylation sites identified by LC-MS/MS following anti-GFP immunoprecipitation of GFP-tagged WT NEK1 from *NEK1* knockout U-2 OS cells, expressed either by transient overexpression or stable doxycycline-inducible expression (Flp-In T-REx system), and analysed using HCD or EThcD fragmentation. All positional numbering is given in NEK1 isoform 3 coordinates (UniProtKB Q5GPYG-3; 1,286 amino acids). Raw positions in the workbook are given in the coordinates of the GFP-NEK1 construct entry used for database searching (MRC_Database_1 entry DU58446: EGFP 1–235; Ser- Gly-Leu-Gly-Ser linker 240–244; NEK1 isoform 3 245–1530); subtract 244 to convert a raw construct position to NEK1 isoform 3 coordinates. Localisation thresholds, per-column definitions and colour-coding are detailed in the workbook’s Methods & Notes sheet; in brief, sites pass where ptmRS ≥85% (or Mascot PTM confidence ≥85% where ptmRS was not computed), with a Mascot ion score >19 required throughout. Where the same peptide backbone appeared with alternative phosphorylated residues, only the highest-confidence localisation is retained per dataset; ambiguous cases are flagged in the Localisation Note column with alternative residue(s) indicated. Phosphosites detected exclusively via multiply-phosphorylated peptides were excluded where the co-phosphorylated residue on the same peptide was independently localised by a singly-phosphorylated species; this avoids double-counting of localisation confidence that cannot be independently attributed to each residue. Sheet guide — OEx HCD WT: all phosphosites detected in the transient overexpression HCD dataset. Stable HCD WT: all phosphosites detected in the stable expression HCD dataset. OEx EThcD WT: all phosphosites detected in the transient overexpression EThcD dataset. Stable EThcD WT: all phosphosites detected in the stable expression EThcD dataset. Common All 4 Datasets: phosphosites detected reproducibly across all four acquisition conditions, with Mascot PTM confidence and ptmRS scores from each dataset shown separately; ten sites met this criterion. PhosphoSitePlus^®^ (NEK1 iso3): all phosphorylation sites currently annotated for human NEK1 isoform 3 in PhosphoSitePlus^®^ [32], with indication of whether each site was detected in the present experimental data. Common All 5 (incl. PSP): Phosphosites common to all four experimental datasets and corroborated by PhosphoSitePlus^®^. Methods & Notes: data processing rules applied throughout, including the multi-phospho peptide curation rule and threshold logic. Summary: overview of the number of phosphosites identified per dataset. ✓, detected; ✗, not detected; —, localisation ambiguous or site not in PhosphoSitePlus^®^. Colour coding (per row): green = detected in both this study and PhosphoSitePlus^®^; yellow = detected in this study, not in PhosphoSitePlus^®^; blue = in PhosphoSitePlus^®^ but not detected; white = not applicable. pThr1096 did not meet the strict four-dataset reproducibility criterion (absent from Stable EThcD) and is retained as a provisional candidate.

**Table S2: Cell-based phosphoproteomic map of kinase-dead NEK1.** Phosphosites detected in KD^D14GA^ GFP-NEK1 across transient overexpression and stable doxycycline-inducible expression datasets, analysed using HCD and EThcD fragmentation. Comparison with the wild-type cell-based phosphosite map in Table S1 and the summary matrix in 5G Figure S5A identifies recurrent sites absent from KD^D14GA^ cells and sites still detected in KD^D14GA^ cells that required recombinant WT/KD^D14GA^ mapping and targeted XIC analysis for further assessment. All positional numbering is given in NEK1 isoform 3 coordinates (UniProtKB Q5GPYG-3; 1,286 amino acids). Raw positions in the workbook are given in the coordinates of the kinase-dead GFP-NEK1 construct entry used for database searching (MRC_Database_1 entry DU80306, GFP-NEK1 D146A: EGFP 1–235; Ser- Gly-Leu-Gly-Ser linker 240–244; NEK1 isoform 3 245–1530), which shares its numbering with the wild-type entry DU58446; subtract 244 to convert a raw construct position to NEK1 isoform 3 coordinates. The D146A substitution abolishes catalytic activity by disrupting the conserved DFG motif required for Mg²⁺-dependent ATP positioning and phosphotransfer; phosphorylation sites detected under this condition therefore represent constitutive or trans-phosphorylation events that do not require NEK1 catalytic activity. Column structure, thresholds and curation rules are as for electronic supplementary material, Table S1 and the workbook Methods & Notes sheet. Phosphosites detected exclusively via multiply-phosphorylated peptides were excluded where the co-phosphorylated residue was independently localised by a singly-phosphorylated species. Sheet guide — Stable HCD KD: stable expression HCD KD^D14GA^ dataset. Stable EThcD KD: stable expression EThcD KD^D14GA^ dataset. OEx HCD KD: transient overexpression HCD KD^D14GA^ dataset. OEx EThcD KD: transient overexpression EThcD KD^D14GA^ dataset. Colour coding — green: site also detected in the wild-type dataset (electronic supplementary material, Table S1) and annotated in PhosphoSitePlus^®^; yellow: in WT dataset, but not in PhosphoSitePlus^®^; blue: in PhosphoSitePlus^®^ but not detected in WT; white: detected only in the KD^D14GA^ condition. Sites detected in both WT and KD^D14GA^ conditions should not be classified as NEK1 autophosphorylation sites on the basis of binary detection alone. Conversely, sites detected in WT, but not KD^D14GA^ cells, represent candidate kinase-activity-associated phosphosites that require quantitative XIC analysis and/or orthogonal biochemical validation for autophosphorylation-site assignment. Autophosphorylation-site assignment in the main text is based on the combined evidence from electronic supplementary material, Tables S1 and S2, recombinant WT/ KD^D14GA^ mapping, XIC quantification where performed (Figure 3B; source data in electronic supplementary material, Table S4), phosphosite mutagenesis and phosphospecific antibody validation.

Table S3: Recombinant NEK1 phosphoproteomic map (wild-type and kinase-dead).

Phosphorylation sites identified by LC-MS/MS on recombinant full-length NEK1 that was expressed in Sf9 insect cells as a GFP-tagged construct, co-expressed with C21ORF2, purified by anti-GFP affinity chromatography, and analysed after RV-3C tag cleavage, comparing WT NEK1 with KD^D14GA^ NEK1. Each condition was analysed in three independent biological replicates using both HCD and EThcD fragmentation. A stringent 3/3 replicate threshold was applied: a site is included only where it was detected in all three replicates of the relevant condition. All positional numbering is given in NEK1 isoform 3 coordinates (UniProtKB Q5GPYG-3; 1,286 amino acids). Raw positions in the workbook are given in the coordinates of the GFP-NEK1 entries of MRC_Database_1 against which the spectra were searched (DU58446 wild-type and DU8030G D14GA: EGFP 1–235; Ser-Gly-Leu-Gly-Ser linker 240–244; NEK1 isoform 3 245–1530; both share this numbering); subtract 244 to convert a raw construct position to NEK1 isoform 3 coordinates. The recombinant protein analysed here was expressed from the baculovirus transfer vectors listed in electronic supplementary material, Table S5, whose tag architecture differs; the raw positions therefore refer to the search-database numbering rather than to that of the expression construct. Column structure and threshold logic are as described for electronic supplementary material, Tables S1 and S2. Phosphosites detected exclusively via multiply-phosphorylated peptides were excluded where the co-phosphorylated residue was independently localised by a singly-phosphorylated species (multi-phospho peptide curation rule; details in Methods & Notes sheet). The 3/3* notation indicates detection in all three replicates by any phospho form (singly or multiply phosphorylated) but in fewer than three replicates by singly-phosphorylated peptide alone. Sheet guide — WT HCD: phosphosites detected 3/3 in WT HCD replicates. WT EThcD: phosphosites detected 3/3 in WT EThcD replicates. D146A HCD: phosphosites detected 3/3 in KD^D14GA^ HCD replicates. D146A EThcD: phosphosites detected 3/3 in KD^D14GA^ EThcD replicates. Common All 4: phosphosites detected 3/3 across all four acquisition conditions. Methods & Notes: data processing rules and curation decisions. Sites absent from KD^D14GA^ datasets, but present in WT datasets, are candidate NEK1 autophosphorylation sites in the recombinant system; sites present in both conditions are constitutive or trans-phosphorylation events acquired from Sf9 insect cell kinases. Comparison with the cell-based WT and KD^D14GA^ maps in electronic supplementary material, Tables S1 and S2, together with XIC quantification where performed (Figure 3B; source data in electronic supplementary material, Table S4), provides orthogonal evidence for autophosphorylation-site assignment in the main text.

**Table S4: Source data for XIC quantification of NEK1 phosphosites** (**Figure 3B**). Wild-type *versus* kinase-dead (WT/KD^D14GA^) peak-area quantification for the candidate phosphosites assessed by XIC. Raw XIC Data: integrated peak areas for each phosphopeptide across WT and KD^D14GA^ conditions and acquisition methods. Calculations: per-site WT/ KD^D14GA^ peak-area ratios across conditions, with minimum and maximum fold-changes. Figure Data (log2): log2-transformed WT/ KD^D14GA^ ratios used to generate Figure 3B. Methods Note: data-processing and normalisation details (peak areas normalised to total ion current).

**Table S5: Key resources and reagents.** Plasmids, antibodies, cell lines and oligonucleotides generated or used in this study, provided as a multi-sheet workbook. Sheet guide — Plasmids – Mammalian: expression constructs for mammalian cell work; Plasmids – CRISPR KO: guide RNA and targeting constructs for *NEK1* knockout generation; Plasmids – Recombinant: constructs for recombinant protein expression in insect cells; Plasmids – Stable Lines: constructs for Flp-In T-REx stable cell line generation; Antibodies: primary and secondary antibodies, including in-house phosphospecific sheep polyclonal antibodies (anti-pSer14, anti-pThr156, anti-pSer418) with antigen sequences and identifiers; Cell Lines: parental and engineered cell lines with sources; Oligonucleotides: primers and guide sequences. Catalogue numbers, internal identifiers, and source details are provided per item where applicable.

**Table S6: Single-variant and gene-burden association statistics for NEK1 in ALS.** Association results for *NEK1* variants from a large-scale exome-wide rare variant study comprising 13,138 ALS cases and 69,775 controls [11]. Variants were tested using Firth logistic regression with the Rare Variant Association Toolkit (RVAT; [78]), with correction for sex, genome-wide burden of synonymous variants, and population structure; a nominal significance threshold of P < 0.05 was applied. Variant nomenclature follows the MANE Select transcript (NM_001155357.3 / ENST00000507142.G), encoding NEK1 isoform 3 (1,286 amino acids). **Sheet 1 (NEK1_sv_full) — single-variant association.** Columns: CHROM, POS (GRCh38 genomic coordinates); ID (dbSNP rsID; “.” if none); REF/ALT (reference and alternate alleles); HGVSc, HGVSp (cDNA- and protein-level nomenclature); domain (annotated protein domain or region); pass_callrate (TRUE/FALSE; see below); caseMAF, ctrlMAF (minor allele frequency in cases and controls); OR, ORCIlower, ORCIupper (odds ratio and 95% confidence interval); *P* (association *P* value). Variants are ordered by ascending *P*. Rows with blank P, OR and MAF fields correspond to variants that were observed but lacked a computable association statistic, because the variant had too few observations (a single observation) to support a valid statistical test. The pass_callrate column indicates whether a variant passes the stringent call-rate quality-control filters used in the exome-wide discovery analysis (filtering to TRUE reproduces the discovery variant set). Variants failing this filter (FALSE) are retained because these thresholds, whilst appropriate for unbiased exome-wide discovery, exclude some established ALS-associated variants; for targeted interrogation of a known ALS gene such as NEK1 these remain informative, consistent with the ALS Gene Curation Expert Panel (GCEP) framework applied in the original study. **Sheet 2 (NEK1_burden) — gene-burden association.** Burden test results (Firth logistic regression) for four variant sets (MAF < 0.05): missense; missense excluding the recurrent p.Arg261His variant; missense + loss-of-function (LOF); and missense + LOF excluding p.Arg261His. Columns: varSetName, bin; nvar; meanCaseScore, meanCtrlScore; OR, ORCIlower, ORCIupper; P. Exclusion of p.Arg261His markedly attenuates the missense-only signal but the missense + LOF burden remains significant, indicating contributions from both the recurrent variant and additional rare deleterious alleles.

